# Genomic Isomerism and Environmental Adaptation in Cyanophages Infecting Freshwater Cyanobacteria

**DOI:** 10.64898/2026.02.14.705889

**Authors:** Yue Shi, Marcus Ziemann, Yang Zhao, Viktoria Reimann, Tao Zhu, Wolfgang R. Hess, Xuefeng Lu

## Abstract

Cyanophages infecting multicellular, heterocyst-forming cyanobacteria play diverse pivotal roles in freshwater ecosystems, yet their infection strategies and genomic adaptability remain poorly understood. Here, we isolated and characterized three *Caudoviricetes* cyanophages (A-Lf14, A-Alj1 and A-Hlh1) specifically infecting the nitrogen-fixing cyanobacterium *Anabaena* sp. PCC 7120. Super-resolution microscopy revealed heterogeneous infection outcomes even among adjacent cells within the same filament, highlighting host-level defense variability. Comparative genomics placed these phages within a cluster that includes the previously isolated phages A-1(L) and N1, and revealed a conserved ∼30 kb invertible genomic region flanked by inverted repeats (IR). This region exists as two stable isomers simultaneously within phage populations, a previously unreported genomic plasticity trait in cyanophages. Consistent with such plasticity, infection kinetics were modulated by light and nitrogen availability, indicating a resource-responsive infection strategy. The novel phages encode an alkaline phosphatase of non-cyanobacterial origin, which was highly upregulated during infection indicating a role in phosphate acquisition in phosphate-limited waters. In contrast, they lack a *tnpB* gene present in A-1(L), which is identical in sequence to five genes in the host genome. We detected protein-coding potential for both strands of *tnpB* and upregulated transcription during infection, consistent with a role in this process. The 347 residues protein encoded on the *tnpB* reverse strand exhibited only limited similarity to other proteins or folding potential, underlining its novelty. Our work illustrates how genomic isomerism, accessory genes, and environmental sensing collectively drive functional diversification in cyanophages, providing insights into phage-host coevolution and the impact of phages on cyanobacterial blooms in dynamic freshwater ecosystems.

## Introduction

Cyanophages are key ecological agents that impact the abundance, diversity, and metabolic outputs of cyanobacteria in freshwater ecosystems^1,2^. By lysing bloom-forming species, they influence nutrient cycling and microbial succession, yet the mechanisms underlying their adaptation to complex, dynamic environments remain poorly understood^2–4^. Compared to cyanophages that target unicellular hosts, this gap is even more pronounced for those infecting multicellular, heterocyst-forming cyanobacteria like *Anabaena*, which are central to nitrogen fixation and certain bloom events^5–8^. Prior to this study, only six cyanophages infecting hosts from the *Anabaena*/*Nostoc* clade had been fully sequenced: A-1(L)^9–13^, N1^11,12^, A-4(L)^11,14,15^, Elbi^16^, YongM^17^, and YongM2. With the exception of A-4(L), these belong to the class *Caudoviricetes*, which encompasses tailed phages with linear, double-stranded DNA genomes^18–20^. The genome of A-1(L) was recently re-sequenced and reannotated, revealing sequence variations compared to the originally deposited version^12,13^. This reannotation also led to the identification of previously unannotated genes, such as those encoding the neck fiber proteins Gp80-Gp82, which form a distinctive bead-chain-like structure^13^. These findings underscore the ongoing need for accurate genomic reference sequences to underpin more detailed mechanistic and molecular studies.

Complementing genomic analysis, high-resolution microscopic and structural techniques are crucial for linking the genetic architecture to functional morphology and infection dynamics^13,21,22^. While advances in cryo-electron microscopy and tomography have revealed conserved viral assembly pathways within unicellular cyanobacteria^21^, these often provide static snapshots^21,23^. For filamentous, heterocyst-forming hosts like *Anabaena*, infection is predicted to involve intricate spatial and temporal coordination across cellular networks^13,24^. However, the application of *in situ* imaging approaches, which are capable of resolving virion transmission, heterogeneous cellular outcomes, and lysis progression, to these complex systems remains notably limited. Overcoming this methodological barrier is therefore essential to uncover how cyanophages navigate, exploit, and ultimately reshape structured multicellular hosts.

Genomic plasticity is a hallmark of viral evolution, enabling rapid adaptation to host defenses and environmental fluctuations. In bacteriophages, site-specific DNA inversion, mediated by dedicated recombinases flipping segments (typically several kilobases) between inverted repeats (IRs), is a classic mechanism for regulating host tropism and gene expression. The classic model for this regulation is the invertible G segment of phage Mu, whose orientation controls the expression of tail-fiber genes and thereby determines host range. Lytic infection strongly selects for one orientation (termed the “+” orientation), whereas induction from lysogeny typically produces a near-balanced mixture of both orientations, directly linking inversion to viral infectivity and lifestyle^25–31^. Although such recombinase-mediated inversion systems have been extensively documented for other bacteriophages (e.g., P1^31–34^, P7^31,33^ and HK022^35^), bacterial pathogens (e.g., *Salmonella*^29–31^), and for the *Saccharomyces cerevisiae* 2-micron plasmid^31,36,37^, they remain scarcely reported in cyanophages. A notable exception is cyanophage Elbi, which carries a ∼30 kb invertible region flanked by a 504 bp IR^16^. Similar IRs of ∼500-700 bp were also noticed in analogous genomic locations in the previously studied cyanophages A-1(L) and N-1. However, it is unclear whether they form part of a functional inversion system^16^. Furthermore, the potential role of other genetic elements such as transposases in shaping phage or host genome architecture and defense remains an open question. This includes TnpB nuclease genes^38–41^, which are found both in the cyanophage A-1(L) and the *Anabaena* host genome^13,42^. Thus, the extent and mechanisms of structural genomic variation in freshwater cyanophages are an intriguing but understudied aspect of virus-host coevolution.

The infection cycle of cyanophages is intimately linked to host physiology, which in turn is profoundly influenced by environmental variables such as light and nutrient availability^7,43–45^. Light fuels host photosynthesis, providing essential energy and carbon for phage replication^45^, while nitrogen availability regulates processes like heterocyst differentiation and overall metabolic activity in filamentous hosts^7,43^. Phosphorus limitation represents another major constraint, particularly in freshwater ecosystems, and triggers sophisticated acclimation responses in both cyanobacterial hosts and their phages^46^. In marine systems, cyanophage infection dynamics are modulated by the host’s phosphorus status and some phages carry auxiliary metabolic genes, including *pstS* encoding the phosphate-binding protein PstS and *phoA*, encoding alkaline phosphatase whose expression is regulated by the host’s phosphate-sensing machinery^47,48^. In freshwater cyanobacteria, the alkaline phosphatase/phosphodiesterase gene *phoD* is widely distributed and plays an important role in scavenging phosphate from organic sources under phosphorus limited conditions^46–50^. Accordingly, the two *phoD* homologs in filamentous *Anabaena* are upregulated during phosphate starvation^46^. However, the presence, role and regulation of similar phosphorus-acquisition genes remains unclear for freshwater cyanophages that infect heterocyst-forming hosts. In fact, it has remained unknown to what extent phosphorus availability shapes phage-host interactions in these environments at all.

To expand the model system for studying filamentous cyanophage-host interactions, we isolated three novel *Caudoviricetes* cyanophages (A-Lf14, A-Alj1 and A-Hlh1) specific to *Anabaena* sp. PCC 7120 from freshwater habitats in Heilongjiang Province, China. Building on prior reports of genomic rearrangements in related phages ^16^, we used comparative genomics to investigate the occurrence and biological relevance of genomic isomerism in these cyanophages. We combined super-resolution microscopy and physiological assays to examine how environmental factors such as light intensity and nitrogen availability modulate infection dynamics, and explored the functional roles of accessory genes *phoD* and *tnpB*. Our findings provide new insights into the multi-layered adaptive strategies of cyanophages, highlighting the dynamic interplay between genomic plasticity, environmental cues, and phage-host coevolution in freshwater ecosystems.

## Results and Discussion

### Isolation and host specificity of novel cyanophages

To isolate novel cyanophages specific to heterocyst-forming cyanobacteria, we collected environmental samples from diverse freshwater habitats in Heilongjiang Province, northeastern China, a region recognized for its rich cyanophage diversity^51–53^. Environmental samples were collected from 36 sites (Supplementary Table 1), spanning a range of aquatic habitats including lakes, ponds, and wetlands, in 2022 and 2023. Filtrates from these samples were screened for lytic activity against the model nitrogen-fixing cyanobacterium *Anabaena* sp. PCC 7120.

From this comprehensive screening, three novel cyanophages, designated A-Lf14, A-Alj1 and A-Hlh1, were isolated based on their ability to cause rapid and complete lysis of the host cultures within 24-48 hours post-infection (Fig. 1a). The lytic activity of these isolates was further confirmed through plaque assays, which revealed clear and distinct lytic halos on lawns of *Anabaena* sp. PCC 7120, demonstrating their infectious nature and confirming the presence of viable virions (Fig. 1b).

**Fig. 1.**
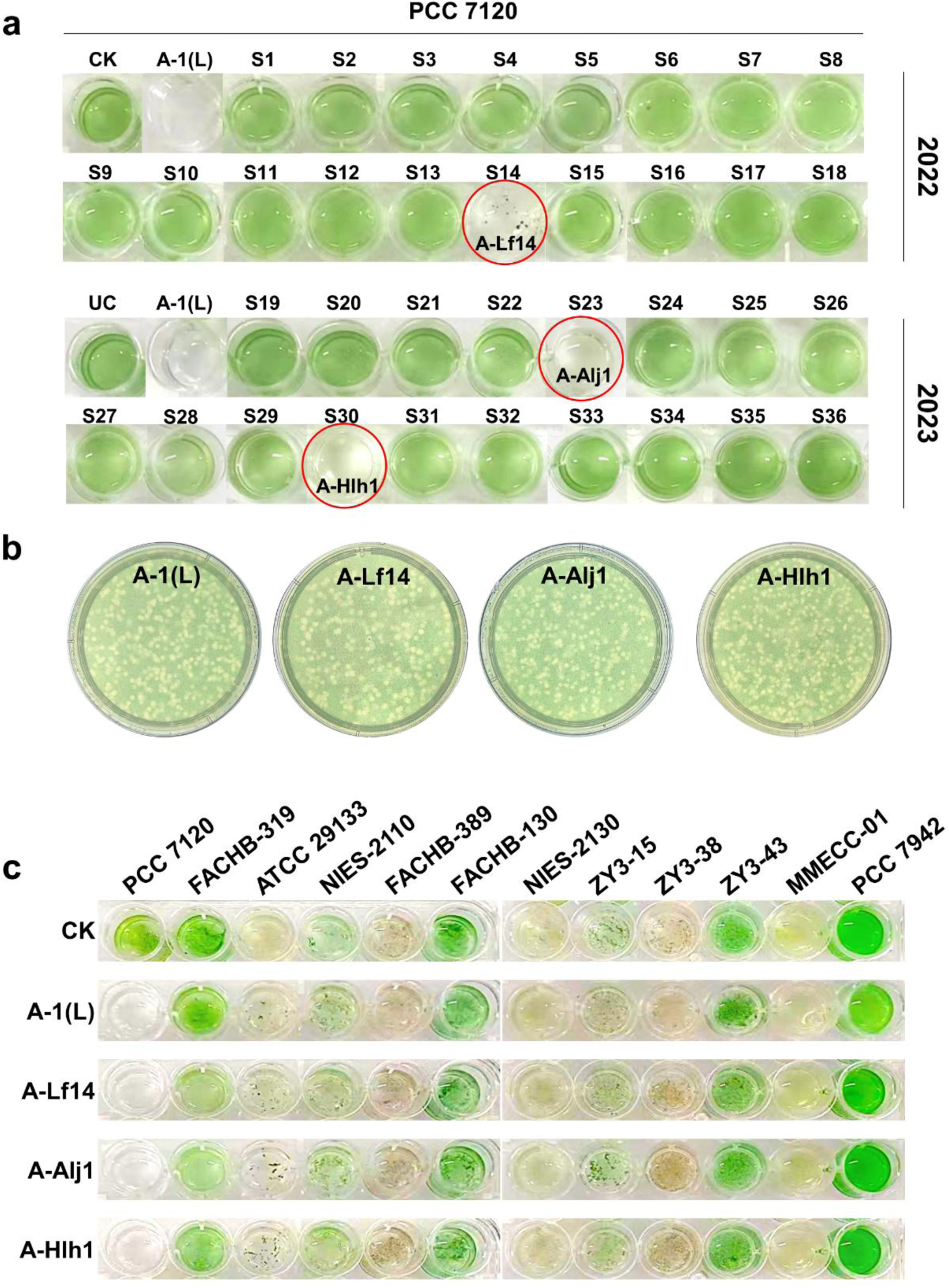
Isolation and host range analysis of novel cyanophages infecting *Anabaena* sp. PCC 7120. (**a**) Primary screening of cyanophage-induced lysis. Environmental filtrates from 36 freshwater sampling sites (S1–S18 collected in 2022; S19–S36 collected in 2023, Supplementary Table 1) were incubated with *Anabaena* sp. PCC 7120 in 24-well plates. The uninfected control (CK) and a positive control infected with the reference cyanophage A-1(L) are shown. Cultures exhibiting complete lysis (chlorosis and clarification) within 24-48 hours were selected for further purification. (**b**) Plaque morphology of cyanophage isolates. Distinct plaque morphologies formed by environmental phage isolates on lawns of *Anabaena* sp. PCC 7120, confirming lytic activity observed in the initial liquid screening. Assays were performed using mid-exponential phase host cultures (OD_730_≈0.5) at a multiplicity of infection (MOI) of 0.01–0.1. Plates were incubated for 72 hours at 30°C under continuous illumination (40 μmol photons m^-2^ s^-1^) using BG-11 medium with a hard agar base (1.5% w/v) and a soft agar overlay (0.6% w/v). (**c**) Host range profiling. Infection assays against 11 phylogenetically diverse cyanobacterial strains, including *Anabaena* sp. PCC 7120, *Anabaena variabilis* FACHB-319, *Nostoc punctiforme* ATCC 29133, *Nostoc* sp. NIES-2110, *Nostoc calcicola* FACHB 389, *Nostoc spongiaeforme* FACHB 130, *Scytonema* sp. NIES-2130, *Nostoc* ZY3-15, *Nostoc* ZY3-38, *Trichormus* sp. ZY3-43, *Limnospira platensis* MMECC-001, *Synechococcus elongatus* PCC 7942, using four *Caudoviricetes* cyanophages [A-1(L), A-Lf14, A-Alj1, A-Hlh1]. Data represent biological triplicates.

Host range analysis against a panel of 11 diverse cyanobacteria revealed that all three novel phages, as well as the reference phage A-1(L), displayed strict specificity for *Anabaena* sp. PCC 7120 (Fig. 1c). No infection was observed against related strains, including *Anabaena variabilis* FACHB-319 and several *Nostoc* species (including strains ZY3-15, ZY3-38, and ZY3-43 isolated from Northeastern China), indicating that infection depended on highly specific host factors not present in the other tested cyanobacteria.

The narrow host range suggests specialized adaptation to *Anabaena* sp. PCC 7120-like hosts, which appear to be present in Chinese freshwater ecosystems despite the strain’s isolation in North America, originally described as *Nostoc muscorum*^9,54^. Moreover, our findings indicate that cyanophages with a similar host range are globally distributed: A-1(L) and A-4(L) were isolated from lakes in the former Soviet Union^10,14^, Elbi from Illinois, U.S.^16^, N-1 from Wisconsin, U.S.^9^, and YongM from lake Dian Chi, China^17^.

The efficient isolation of these phages through direct host screening and sequential enrichment indicates they are well-adapted to laboratory cultivation, making them suitable models for further molecular and mechanistic studies. Their specificity also highlights their potential role in selectively regulating host populations in natural environments. Finally, they constitute a valuable addition to the limited number of isolated phages infecting heterocyst-forming cyanobacteria.

### Morphological characterization and visualization of infection

To close the critical knowledge gap in visualizing cyanophage infection of structured multicellular hosts, we applied a multi-modal imaging approach for characterization of virion morphology and dissection of the infection process. Transmission electron microscopy (TEM) of negatively stained particles confirmed that A-Lf14, A-Alj1, and A-Hlh1 all shared the canonical *Caudoviricetes* morphology of an icosahedral capsid and a contractile tail (Fig. 2a). Notably, all three phages displayed elaborate neck fibers (Fig. 2a, yellow arrows), a feature structurally reminiscent of the bead-chain-like neck fibers formed by Gp80–Gp82 in cyanophage A-1(L)^13^. However, these newly isolated cyanophages possess neck fibers that are substantially longer than those of A-1(L)^13^, a structural difference that likely underpins distinct host-interaction strategies during initial attachment.

**Fig. 2.**
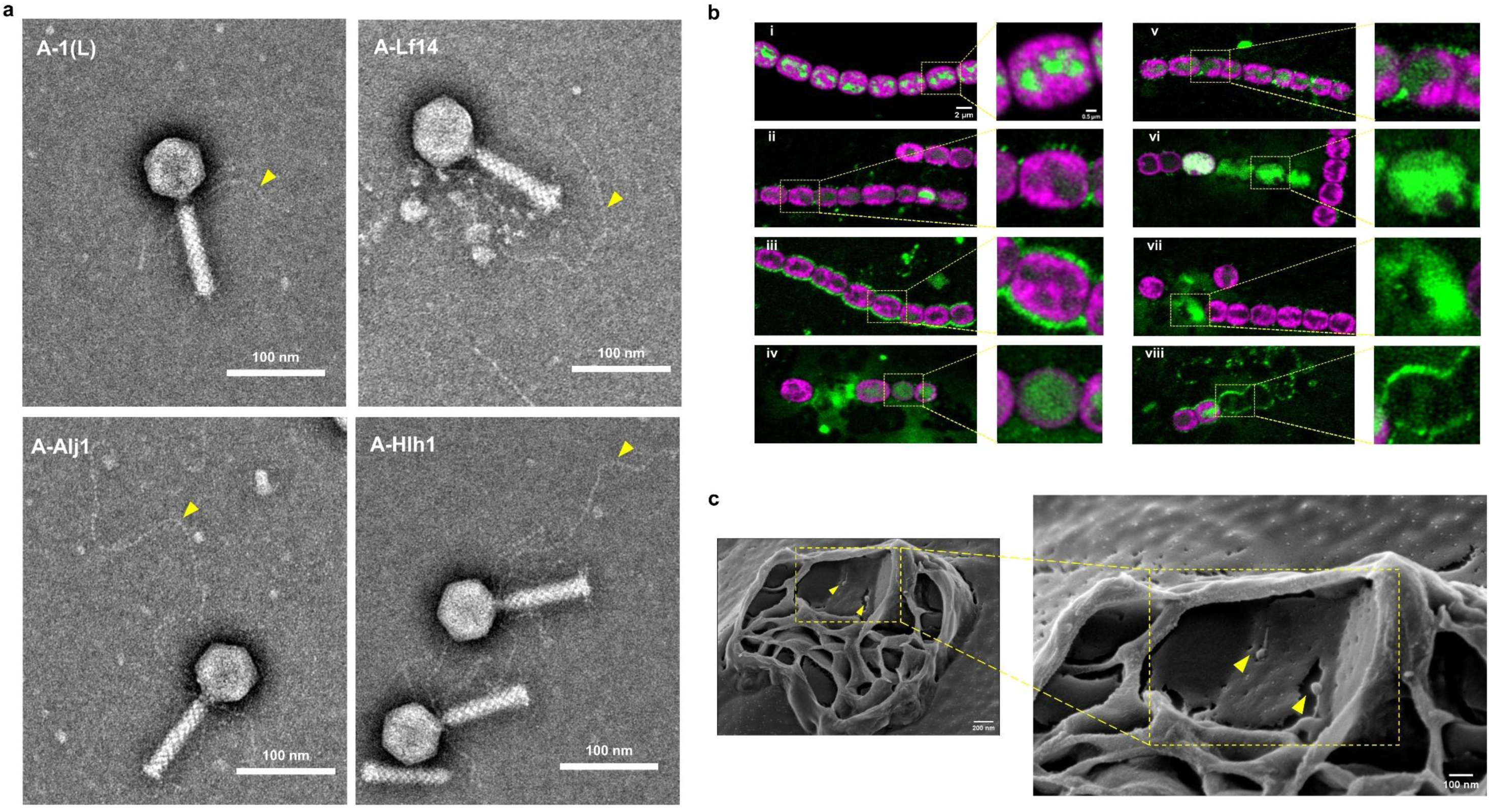
Morphological characterization of cyanophage virions and infection progression in *Anabaena* sp. PCC 7120. (**a**) TEM of negatively stained cyanophage particles. Virions of A-1(L), A-Lf14, A-Alj1, and A-Hlh1, stained with 3% uranyl acetate, display characteristic *Caudoviricetes* (Myovirus) morphology, with icosahedral capsids (64 ± 2 nm in diameter, n = 20), contractile tails (95 ± 5 nm in length, n = 20). Notably, the neck fibers (yellow arrows) in A-1(L) are substantially shorter than those observed in the three novel isolates. (**b**) Visualization of cyanophage A-Lf14 infection by confocal fluorescence microscopy. Samples containing uninfected or phage-infected *Anabaena* filaments were stained with DAPI (pseudocolored green; ex/em 401/422 nm) and imaged using a Zeiss LSM 880 microscope with Airyscan detection. Host phycobilisome autofluorescence is shown in magenta (ex/em 548/561 nm). The overlay images depict key stages of the infection cycle: (i) uninfected control filament; (ii–iii) phage adsorption and accumulation on the cell surface; (iv) intracellular virion assembly; (v-vii) release and propagation; (viii) phage particles associated with empty cellular sheaths. (**c**) Intracellular virion assembly observed by cryo-scanning electron microscopy (cryo-SEM). *Anabaena* sp. PCC 7120 cells infected with A-1(L) were plunge-frozen at 8 hours post-infection. Arrows indicate mature phage particles within the host cytoplasm, exhibiting typical capsids and tails consistent with the *Caudoviricetes*. The corresponding enlarged view of the boxed area is presented on the right, highlighting the assembled virions. Imaging was performed on a Zeiss Ultra Plus HR-SEM at 2 kV with a stage temperature of −145 °C.

Airyscan super-resolution confocal microscopy was used to visualize the infection process, focusing on cyanophage A-Lf14 as a model. To capture distinct morphological stages spanning from phage adsorption to host lysis we used 4’,6-diamidino-2-phenylindole (DAPI) as a DNA stain (Fig. 2b). Relative to uninfected control filaments (Fig. 2b, panel i), our imaging analysis delineated key steps in this progression: phages accumulating on the filament surface (Fig. 2b, panels ii-iii), followed by a phase marked by intense intracellular DAPI signals indicative of virion assembly (Fig. 2b, panel iv), and culminating in the apparent release of progeny and phage particles associated with cellular debris (Fig. 2b, panels v-viii).

Three-dimensional reconstruction of infected filaments provided deeper spatial insights into host-phage interactions and revealed population heterogeneity. Super-resolution Z-stacks revealed variable infection patterns: from peripheral focal accumulations during initial phage adsorption and DNA entry (Extended Data Fig. 1, Panel Infection 1), to dispersed signals indicating host genome disruption and likely progeny virion release. This process culminated in the formation of cellular debris enriched for phage and/or host nucleic acids (Extended Data Fig. 1, Panel Infection 2). A volumetric rendering of a representative filament further resolved spatially distinct scenarios within a single host, including surface-adsorbed phages, potential cell-to-cell spread, and clusters of intracellular virions (Supplementary Video 1), offering unprecedented detail of infection architecture within a filamentous cyanobacterium.

To obtain ultrastructural corroboration at nanometer resolution, we performed cryo-scanning electron microscopy (cryo-SEM). Imaging of cell filaments infected with A-1(L) at 8 hours post-infection provided direct evidence of mature virions assembled within the host cytoplasm (Fig. 2c). Critically, cryo-SEM captured the striking asynchrony of phage-induced lysis, where two neighboring cells within the same filament exhibited vastly different degrees of disruption (Extended Data Fig. 2a). A series of single-cell views detailed a continuum of structural damage, from early interior alterations to complete cellular disintegration (Extended Data Fig. 2b-i), directly visualizing the single-cell heterogeneity of infection outcomes in a multicellular context. This integrated imaging framework spanning detailed virion architecture, life cell confocal tracking, multi-dimensional fluorescence mapping, and high-resolution cryo-electron microscopy, validates a highly productive and asynchronous lytic cycle. The efficient intracellular virion assembly visualized here is consistent with the high burst sizes reported for related phages^11,15,55^. More importantly, it provides a comprehensive visualization strategy to dissect infection dynamics and host-level variability in complex, structured multicellular systems.

### Whole-genome sequencing, taxonomic characterization *and c*omparative genomic analysis of A-Lf14, A-Alj1 and A-Hlh1

To contextualize the newly isolated cyanophages within the broader landscape of cyanophage diversity and evolution, we performed high-quality whole-genome sequencing of A-Lf14, A-Alj1 and A-Hlh1, and re-sequenced the genome of A-1(L). These data, combined with genomes of the five most closely related phages (Supplementary Table 2), enabled a comprehensive comparative analysis that not only confirmed their taxonomic placement, but also revealed novel genomic features underlying their adaptation. All three isolates possess double-stranded DNA genomes, with sizes of 70,561 bp (A-Lf14), 70,112 bp (A-Alj1), and 71,024 bp (A-Hlh1). These lengths are similar to that of the well-characterized cyanophage A-1(L)^12,13^ (Supplementary Table 2), further supporting their classification within a cohesive morphological and genomic group. To elucidate their evolutionary origins and genomic conservation patterns, we leveraged these sequences alongside with publicly available cyanophage genomes to reconstruct phylogenetic relationships and assess gene-content variation within this ecologically relevant viral lineage.

Phylogenomic analysis was conducted using a whole-proteome approach implemented in the program Viptree^56^. A total of 280 complete cyanophage genomes retrieved from GenBank and the Virus-Host Database^57,58^ were included, alongside the three new isolates (Fig. 3a; see Supplementary Table 3 for accession list). This analysis placed A-Lf14, A-Alj1 and A-Hlh1 within a well-defined clade consisting of six previously described *Caudoviricetes* phages that infect *Anabaena* or related cyanobacterial hosts: *Nostoc*/*Anabaena* phages A-1(L), N1, YongM, YongM2, and Elbi, as well as *Dolichospermum* phage Dfl-JY23. This robust phylogenetic clustering, consistent with the shared *Caudoviricetes* morphology observed by TEM (Fig. 2a), underscores the evolutionary coherence of this group and provides a firm framework for our subsequent genomic comparisons.

**Fig. 3.**
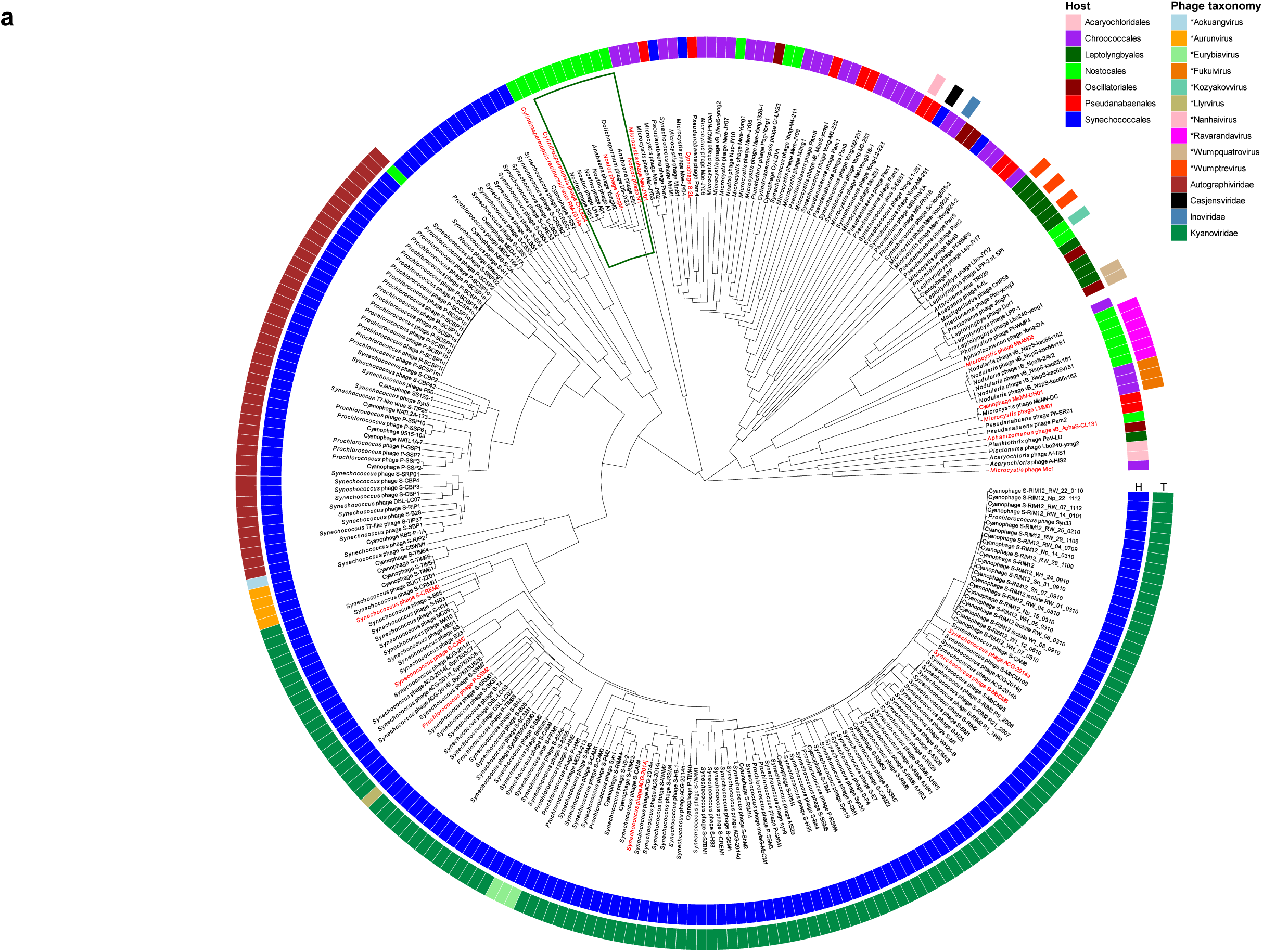
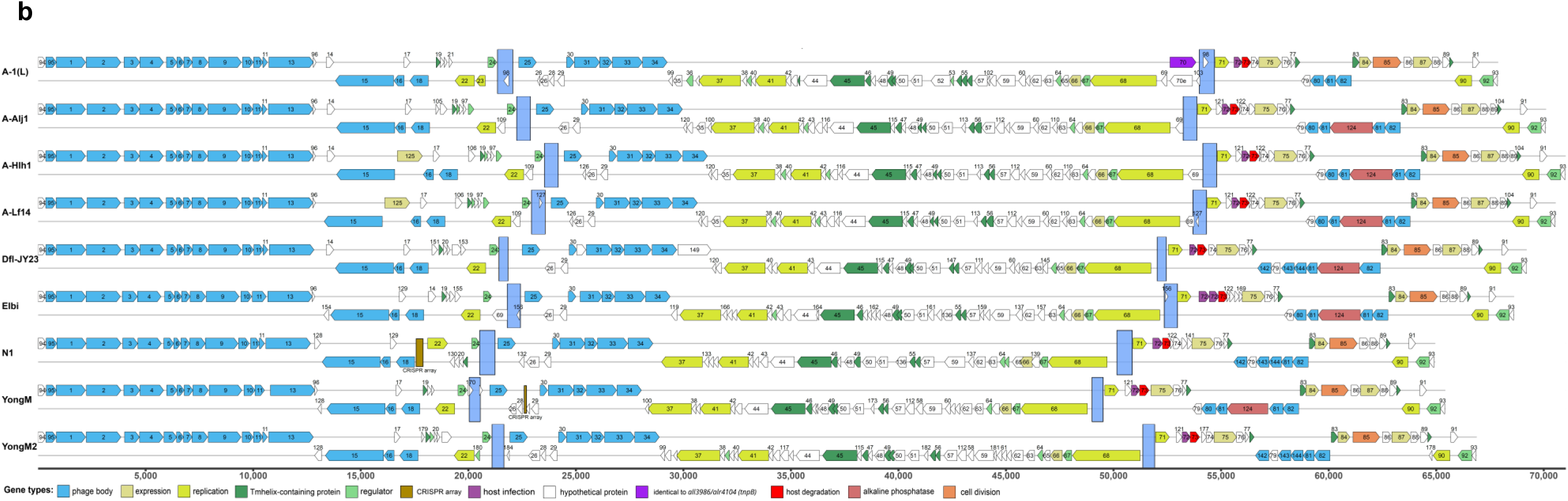
Phylogenomic and comparative genomic analysis of IR-containing cyanophages. (**a**) Phylogenomic analysis of 280 cyanophage whole-proteomes using Viptree ^56^. The tree is presented at logarithmic scale, host (H) and phage taxonomy (T) were added to the picture. Phages encoding CRISPR arrays are highlighted in red. The clade containing *Anabaena* phage A-1(L) and related isolates is boxed in green. (**b**) Comparative genomic analysis of nine cyanophages containing a conserved invertible genomic element (blue rectangle). Linear genome maps are shown, with genes colored according to predicted function and labeled with their corresponding gene cluster numbers. For better comparison, all genomes were re-drawn, with the start codon of the gene cluster 94 at the left terminus and all invertible regions oriented consistently. See Extended Data Fig. 3 for an alignment of the IR sequences flanking the invertible regions.

Comparative genomic analysis of the nine phages in this clade revealed a core set of 62 universally conserved gene clusters, alongside numerous lineage- and isolate-specific genes (Fig. 3b). In total, 181 gene clusters were identified, of which 82 could be assigned predicted functions or protein domains through homology searches and curated references^12,13^. These included 28 gene clusters encoding structural phage proteins, 13 transmembrane-domains-containing proteins, 15 regulatory factors, and 9 involved in DNA replication (Supplementary Table 4). Only 69 clusters are conserved among all nine phages and 56 exist exclusively in one phage (Supplementary Table 4). Most of the 56 isolate-specific clusters corresponded to short sequences (<300 nt) lacking detectable similarity to known genes and could represent rapidly evolving genes such as small anti-CRISPR proteins, or possibly, annotation artifacts. An analysis for anti-CRISPR proteins, based on the algorithms LSTM, GRU and LINEAR^59^ indicated proteins in the clusters P039, P121, P160 as possible candidates (Supplementary Table 4). Detailed information on all 943 proteins annotated in the 9 genomes is included in a separate functional inventory (Supplementary Table 5).

### Identification of a conserved invertible genomic element

Comparative genomic alignment across the nine related cyanophages unveiled a striking and conserved structural feature: a large, syntenic region flanked by inverted repeats (IRs) ranging from 400 to 700 nt in length (Fig. 3b). Alignment of these IR sequences shows short conserved motifs, but these are not shared among all phages (Extended Data Fig. 3). Despite the sequence divergence, the consistent presence of these IRs in analogous genomic locations suggests that they serve as recognition sites for a conserved rearrangement mechanism.

Remarkably, whole-genome alignments revealed that the IR-bounded segment occurred in opposite orientations across different isolates: for example, A-1(L), A-Hlh1 and N1 shared one orientation, whereas A-Alj1, A-Lf14 and Elbi exhibited the reverse orientation. This lack of correlation between inversion state and phylogenetic relatedness indicates that the direction of the invertible element is not a fixed trait and is unlikely to result from a single ancient evolutionary event. If the inversion were phylogenetically static, closely related phages would be expected to share the same orientation; the observed mosaic distribution instead strongly implies that inversion occurs recurrently within lineages, likely contributing to genomic plasticity.

To facilitate direct visual comparison, all genomes, which are canonically linear, double-stranded DNA molecules, were re-drawn with the start codon of the conserved cluster 94 positioned at the left terminus and the invertible region uniformly oriented (Fig. 3b). The presence of both orientations within closely related isolates raised the key question of whether these inversions represent genuine biological polymorphisms or potential assembly artifacts arising from the repetitive IR sequences, prompting us to pursue experimental validation (see next section). Overall, the identification of this conserved yet orientation-labile genomic element highlights a previously overlooked dimension of structural plasticity in freshwater cyanophages, setting the stage for investigating its mechanistic basis and functional implications in virus-host adaptation.

### Experimental validation of genomic isomerism in cyanophage populations

To experimentally determine whether the observed IR-flanked genomic inversion represents a true biological polymorphism or a technical artifact, we first visualized the genomic organization of the four cyanophages. Comparative alignment using GBKviz clearly illustrated the syntenic region flanked by IRs, with the orientation of the intervening segment reversed in different phages (Fig. 4a). This inversion was observed between A-1(L) and A-Hlh1 (one orientation) and A-Alj1 and A-Lf14 (opposite orientation), supporting the bioinformatic prediction of genomic isomerism.

**Fig. 4.**
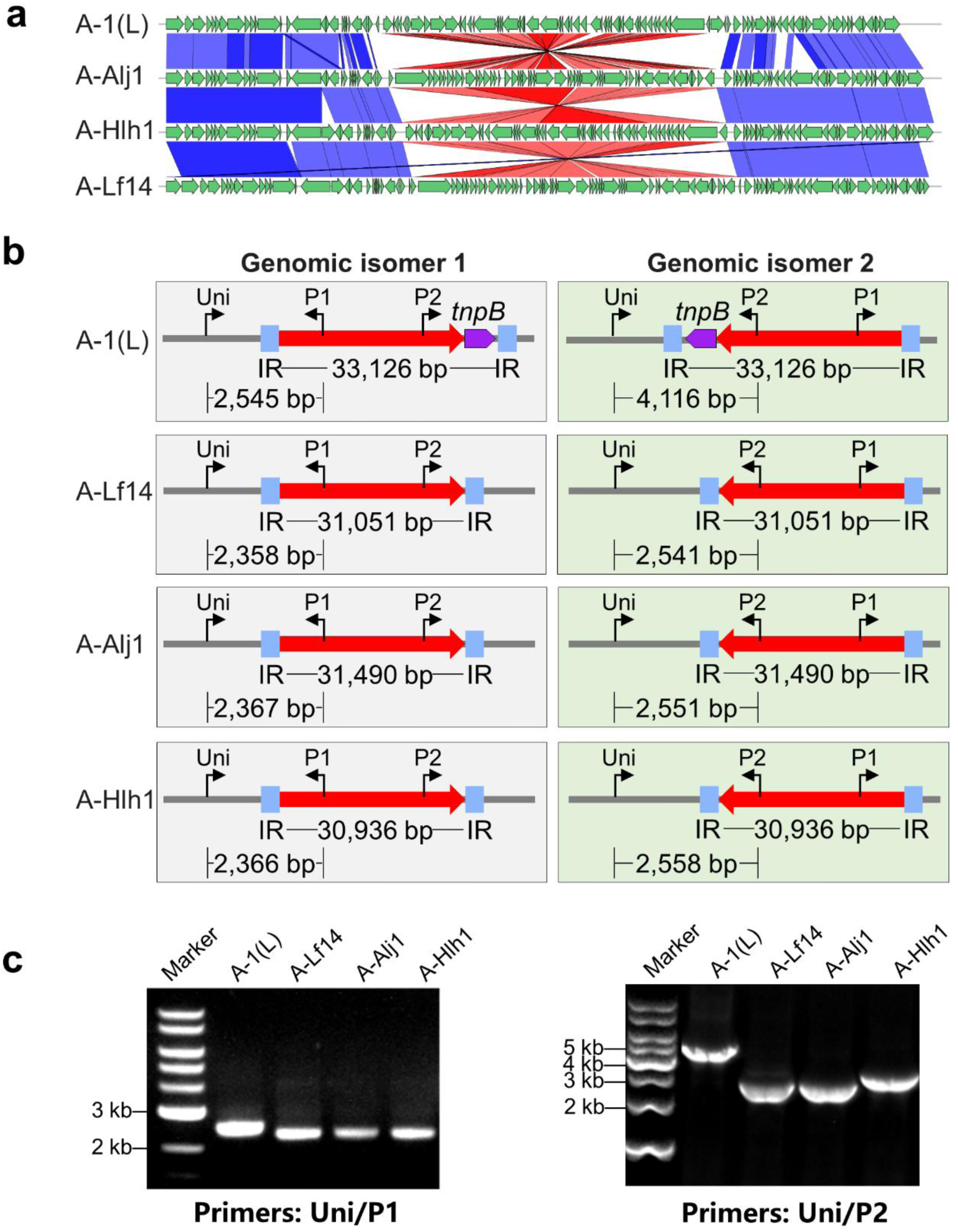
Reversible orientation isomers of a flexible genetic module in *Anabaena*-infecting *Caudoviricetes* cyanophages. (**a**) Comparative genomic visualization of the four *Anabaena* sp. PCC 7120-infecting cyanophages. Genomic comparisons were performed using GBKviz. The alignment relationships between the conserved flanking regions across the four phages are connected and shown in blue, while those between the internal invertible modules are connected and highlighted in red. (**b**) Schematic of the two genomic isomers and diagnostic PCR strategy. The diagram illustrates the two orientation isomers (Isomer 1 and 2) and shows the specific structures of phages A-Lf14, A-Alj1, and A-Hlh1. Inverted repeats (IRs) that flank the module are marked. To distinguish the isomers, a unique primer ‘Uni’ anchored in the flanking stable region was paired with outward-facing primers P1 or P2 located within the conserved invertible module, enabling specific amplification of each isomer. The predicted sizes (in bp) of the resulting Uni/P1 and Uni/P2 amplicons are indicated for each phage, and the length of the invertible fragment itself is annotated. For reference in later comparative analysis, the transposase gene *tnpB*, which located merely inside the invertible module and adjacent to the IR in phage A-1(L), is also labeled in the schematic. (**c**) Experimental validation of both isomers by PCR. Agarose gel electrophoresis of PCR products from the four cyanophage isolates using diagnostic primer pairs Uni/P1 (left panel) and Uni/P2 (right panel). Molecular size markers are indicated in lane M. The observed band sizes correspond to the predictions in panel (**b**), confirming the coexistence of both isomeric configurations in each phage culture.

To directly test for the coexistence of both isomers within phage populations, we designed a diagnostic PCR assay. Primer pairs Uni/P1 and Uni/P2 were used to specifically amplify the two isomeric configurations (Fig. 4b). PCR products of expected sizes were obtained from all four cyanophage isolates with both primer sets, confirming that each culture contained viral particles harboring the IR-bounded segment in both orientations (Fig. 4b and c). The amplicon lengths precisely matched the predicted sizes, and Sanger sequencing of these products confirmed their exact nucleotide correspondence to the two isomeric configurations (Fig. 4b and c; Extended Data Fig. 4), providing molecular validation of the inversion.

We also sought to exclude the possibility that these isomers arose as artifacts during *de novo* genome assembly. To this end, we mapped raw Illumina sequencing reads independently to both isomeric reference sequences. This orthogonal analysis revealed unambiguous and abundant read coverage across each orientation for all phages (Extended Data Fig. 5), confirming at the population sequence level that both isoforms genuinely coexisted within the same viral stock.

A broader search across publicly available cyanophage genomes identified structurally similar IR-flanked regions in five additional phages, which infect taxonomically distant hosts such as *Cylindrospermopsis*, *Synechococcus*, and *Microcystis* (Supplementary Table 6). The presence of analogous invertible elements in phylogenetically distinct cyanophages suggests that this genomic architecture is conserved within a limited subset of tailed phages and may play a role beyond the *Anabaena*-infecting group.

The observed coexistence of both genomic isomers within propagated populations raises the question of the underlying mechanism. While the inversion could be mediated by a site-specific recombinase, the natural genomic fluidity of many *Caudoviricetes* phages provides an alternative explanation. For cyanophages infecting *Anabaena*, such as A-1(L), N-1, and others, replication proceeds via circular intermediates and concatemers ^60–64^, with genomes packaged by a headful mechanism that generates circularly permuted and terminally redundant ends^20,62–64^. This intrinsic recombination-prone architecture, characterized by dynamic terminal repeats and heterogeneous genome ends, may itself facilitate or stabilize large-scale genomic rearrangements, including the maintenance of segmental isomers, even in the absence of a dedicated recombinase^65^. It is worth noting that in the classic phage Mu model, the orientation of the invertible G segment is influenced by the viral lifestyle, with lysogenic induction often yielding a balanced mixture of both isomers^25–31^. Although it remains unclear whether the cyanophages studied here have undergone lysogeny, a similar lifestyle-associated mechanism could potentially influence the isomer ratios observed in our propagated populations.

Thus, combined PCR, sequencing, and read-mapping data provide robust multi-method evidence that the IR-bounded genomic segment exists as two stable, reversible isomers within propagated cyanophage populations. This genomic plasticity likely represents a biologically functional trait, possibly contributing to phenotypic diversity or adaptation in infection dynamics, although its precise mechanistic role remains to be elucidated.

### Environmental regulation of cyanophage infection dynamics

The infection cycle of cyanophages is profoundly influenced by the physiological state of their cyanobacterial hosts, which in turn is modulated by key environmental factors. Light plays a central role in all stages of the cyanophage life cycle, from adsorption to replication. Furthermore, in filamentous nitrogen-fixing cyanobacteria such as *Anabaena* sp. PCC 7120, nitrogen availability critically regulates heterocyst development and has been shown to shape the evolution of phage resistance ^7,8^. Therefore, we examined how variations in light intensity and nitrogen supply affect the infection dynamics of our isolated cyanophages.

Our results indicated that light intensity did not function as a simple binary switch for infection but rather acted as a fine-tuner of its kinetics (Fig. 5a). Under standard light conditions, all four phages induced rapid and complete lysis of the host within 24 h post-infection. When infection was initiated in darkness, the lytic process was significantly delayed during the first 24 h; however, full lysis was still achieved by 48 h, suggesting that phages can mobilize stored host energy reserves or sustain replication under severely reduced metabolic activity. Intriguingly, exposure to high light did not accelerate lysis but instead slightly decelerated the initial decrease in optical density for most phages, though complete lysis still occurred within 48 h. This may reflect transient photoinhibitory stress on the host, temporarily slowing the metabolic pathways co-opted by the phage. A distinct pattern was observed for phage A-1(L): its lysis was slower under standard light but comparatively faster in darkness after 36 h, pointing to potential genetic adaptations for resource utilization under energy-limited conditions.

**Fig. 5.**
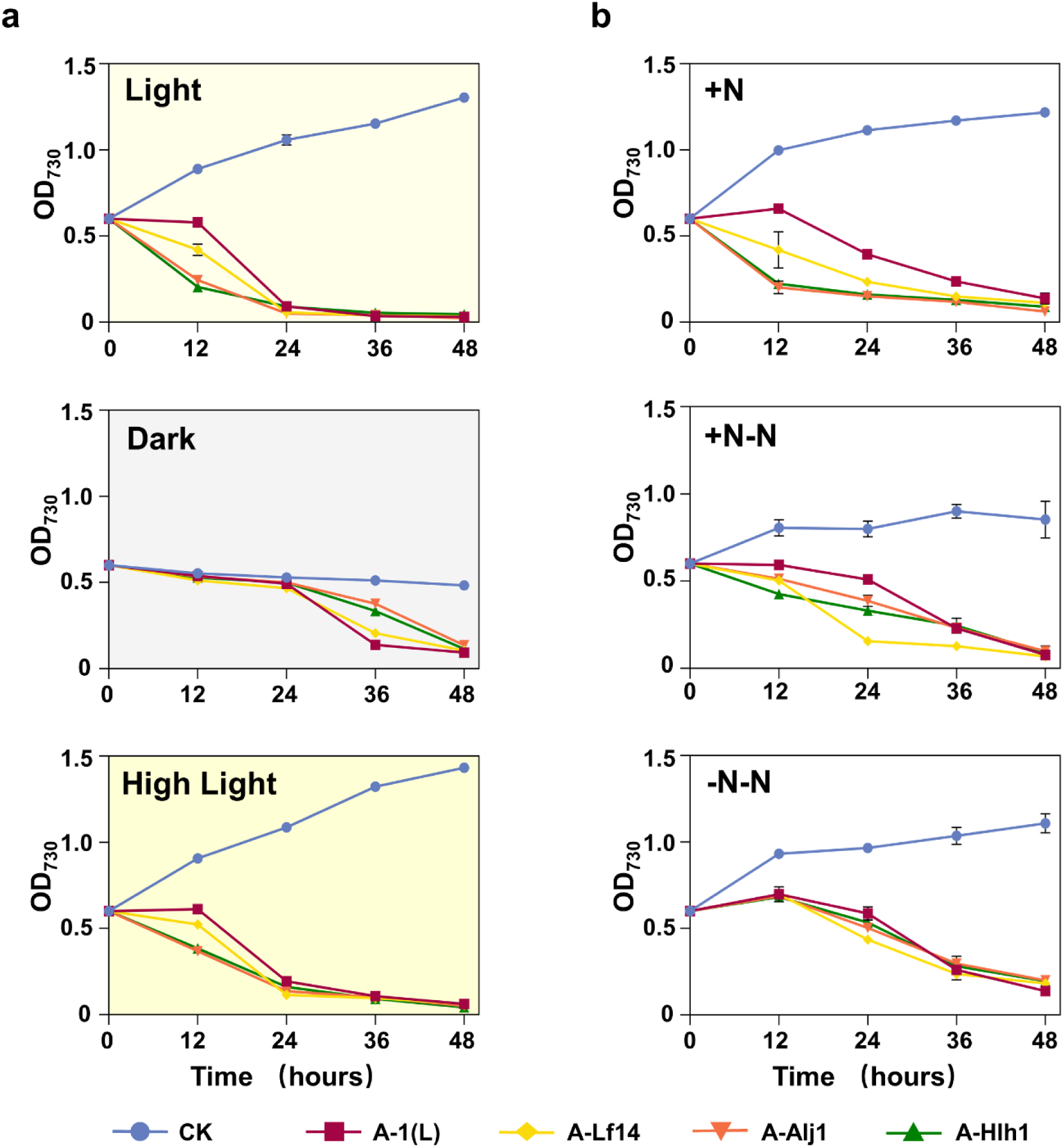
Environmental regulation of cyanophage infection dynamics in *Anabaena* sp. PCC 7120. (**a**) Light intensity modulates lytic kinetics. Growth curves for cyanophages A-1(L), A-Lf14, A-Alj1, and A-Hlh1 were determined under varying light conditions: standard light (40 μmol photons m^-2^ s^-1^), darkness, and high light (150 μmol photons m^-2^ s^-1^). Uninfected controls are included for each condition (CK). (**b**) Nitrogen availability constrains infection progression. Infection profiles were assessed under different nitrogen regimes: nitrogen-replete conditions (+N), nitrogen-deprived infection in nitrogen-replete cultured cells (+N-N), and nitrogen-deprived infection in nitrogen-starved cells (-N-N). Uninfected controls for each nitrogen treatment are presented (CK). Data represent means ± SD of three biological replicates. Cultures were grown in BG-11 medium under standard conditions unless otherwise specified, and infection was monitored by measuring culture turbidity at OD_730_ over a 48-hour period.

A similar kinetic modulation was observed in response to nitrogen availability (Fig. 5b). In nitrogen-replete medium (+N), all phages induced rapid and robust host lysis. In contrast, when nitrogen was removed at the time of infection (+N-N) or in cultures pre-starved of nitrogen (-N-N), the lytic cycle was markedly attenuated. Importantly, lysis was ultimately completed in both nitrogen-depleted treatments by 48 h, indicating that while accessible nitrogen pools are crucial for rapid phage propagation, the phages can eventually scavenge nitrogen from pre-existing cellular macromolecules to support replication.

Together, these findings demonstrate that the pace of cyanophage infection is highly plastic and tightly coupled to the metabolic state of the host. Rather than being an all-or-nothing response, the lytic outcome is achieved through kinetic delays that reflect the phage’s capacity to adapt to and exploit a resource-limited intracellular environment. This environmental responsiveness underscores the ecological flexibility of cyanophages and provides a physiological context for interpreting the genomic and structural features described earlier.

### Comparative genomic analysis reveals functional differentiation in cyanophage isolates

The phenotypic plasticity prompted us to search for underlying genomic determinants that might confer adaptive advantages under specific environmental conditions. Our high-fidelity sequencing and systematic validation efforts have culminated in a definitive genome sequence for the cyanophage A-1(L). Comparative analysis of the three available high-throughput sequencing datasets identified 12 genomic variants^12,13^ (Supplementary Table 7). Through comprehensive Sanger sequencing, we resolved 9 of these variants to yield a single, high-confidence sequence, while confirming 3 sites as genuine polymorphisms within the viral population (Supplementary Table 7). This refined genome provides a relevant reference for future functional studies of this model cyanophage.

Expanding this comparative genomics framework by including the closely related cyanophages A-Lf14, A-Alj1, and A-Hlh1 uncovered a suite of accessory genes that were differentially present or absent across the phage genomes, including putative transcription factors, antirepressors, restriction enzymes, and distinct neck - fiber variants (Supplementary Table 5; Fig. 3b). Among these variable loci, two genes stood out due to their contrasting distribution and direct relevance to environmental adaptation and genomic plasticity: *phoD* and *tnpB*. The *phoD* gene (cluster P124 in Supplementary Table 4), encoding an alkaline phosphatase, was present in the three novel isolates (A-Lf14, A-Alj1, and A-Hlh1) but absent in A-1(L), whereas the *tnpB* gene (cluster P70 in Supplementary Table 4), encoding a transposase-associated protein, showed the opposite pattern (Supplementary Table 8). This reciprocal distribution highlights the divergent evolutionary trajectories of these closely related phages and led us to focus subsequently on these two environmentally responsive genetic modules. The phage-encoded PhoD proteins exhibited low amino acid identity (<30%) to the two host *Anabaena* sp. PCC 7120 homologs, PhoD1 (Alr2234) and PhoD2 (Alr4976)^49^, but showed higher sequence similarity to PhoD proteins from other bacteria, such as the Verrucomicrobiia. This pattern suggests acquisition from a non-cyanobacterial source, potentially expanding the phage’s metabolic capabilities. Indeed, our assays showed that while A-Hlh1 (carrying *phoD*) and A-1(L) (lacking *phoD*) both infected the host efficiently under phosphate-replete conditions, only A-Hlh1 infection was strongly inhibited under phosphate depletion (Fig. 6a). Transcript analysis revealed that under both phosphate-replete and phosphate-depleted conditions, the expression of *phoD2* was significantly higher than that of *phoD1*, indicating that PhoD2 is the predominant alkaline phosphatase in the host. Although phosphate depletion induced a greater fold change in *phoD1* expression, its absolute transcript level remained substantially lower than that of *phoD2* (Fig. 6b). During infection, the phage-borne *phoD* genes in all three novel isolates were dramatically upregulated, exceeding host *phoD* transcript levels by nearly two orders of magnitude, indicating a conserved and dominant role for phage-derived alkaline phosphatase (Fig. 6c). This suggests that the phage *phoD* is deployed to actively manipulate host phosphate metabolism, a strategy that becomes critical when environmental phosphate is scarce. Furthermore, the *phoD* gene was consistently located adjacent to neck fiber (*nf*) structural genes, forming a conserved genomic module. However, expression analyses across A-Lf14, A-Alj1, and A-Hlh1 revealed independent transcription patterns for *phoD* and the adjacent *nf* genes (Fig. 6d, e), indicating that *phoD* is not part of the structural gene operon and its expression is likely subject to separate, condition-dependent regulation. The exceptionally high expression of the *nf2* gene in A-Lf14, A-Alj1, and A-Hlh1 is consistent with previous structural data on A-1(L), in which the homologous *gp81* gene encodes a homotrimeric protein forming 14 repetitive units in the neck fiber assembly^13^. Notably, sequence divergence within these *nf* genes likely underlies the substantial difference in neck fiber length observed between the newly isolated phages and A-1(L) (Fig. 2a), providing a genetic basis for their distinct host-interaction structures.

**Fig. 6.**
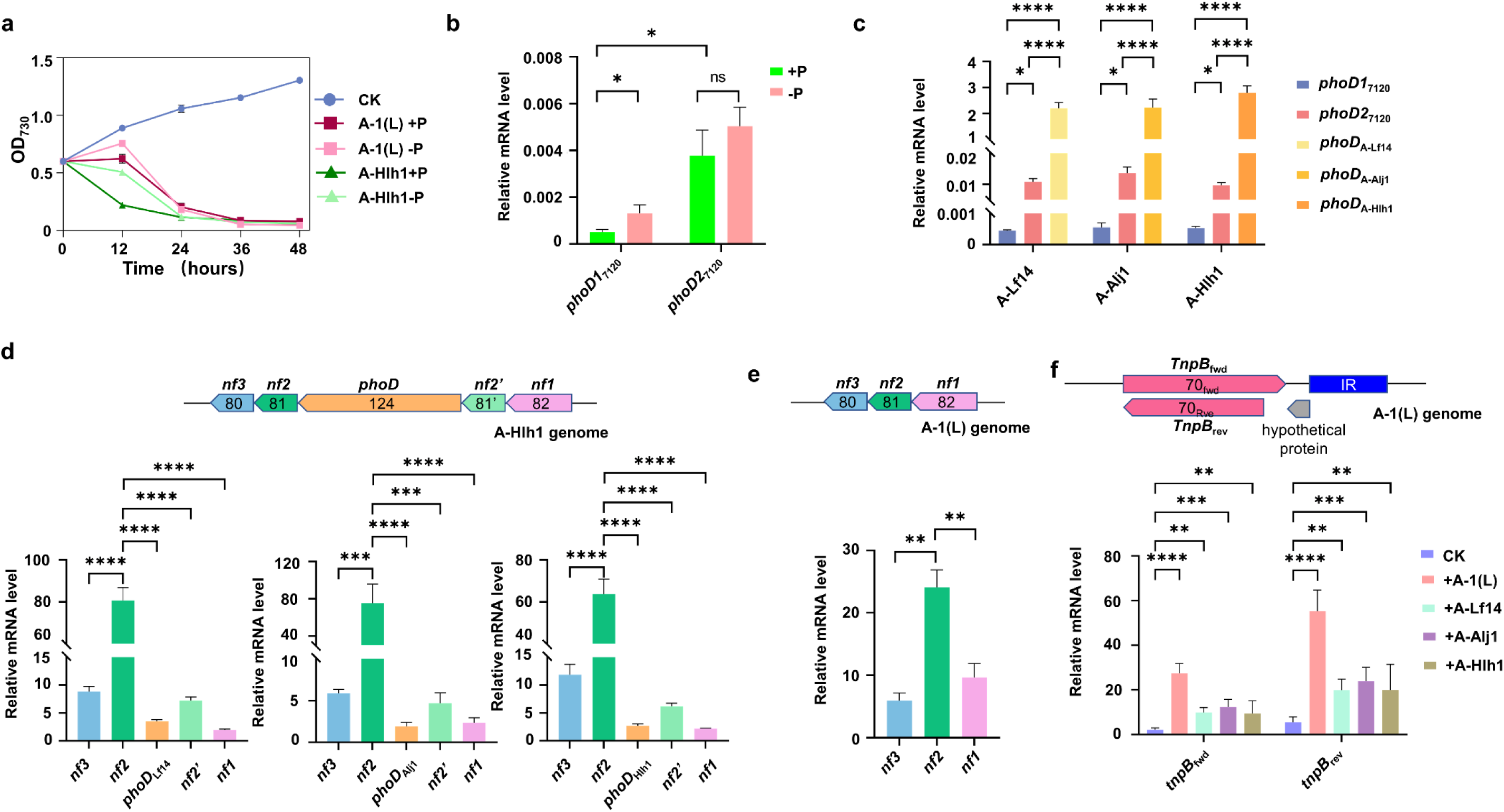
Functional analysis of phage-encoded accessory genes and their expression dynamics during infection. (**a**) Phosphate-dependent infection kinetics. Growth curves of cyanophages A-1(L) and A-Hlh1 under phosphate-replete (+P) and phosphate-depleted (-P) conditions over 48 hours. The infectivity of A-Hlh1, which carries a *phoD* gene, was strongly inhibited under -P conditions, while A-1(L), which lacks *phoD*, remained relatively unaffected. (**b**) Differential induction of host alkaline phosphatase genes under phosphate stress. Transcript levels of *phoD1* (*alr2234*) and *phoD2* (*alr4976*) in uninfected *Anabaena* sp. PCC 7120. Both genes showed basal expression under +P conditions, with *phoD2*_7120_ levels being higher than *phoD1*_7120_. Phosphate depletion induced the expression of both genes, with a more pronounced fold induction observed for *phoD1*_7120_. (**c**) Dominant expression of phage-encoded *phoD* during infection. Relative transcript abundance of the phage-borne *phoD* genes (A-Lf14, A-Alj1 and A-Hlh1) and host *phoD1*_7120_ and *phoD2*_7120_ genes during infection. In all three *phoD*-carrying phages, the phage-encoded *phoD* was expressed at levels nearly two orders of magnitude higher than that of the host *phoD* genes, indicating a conserved and dominant role of phage-derived alkaline phosphatase. (**d**) Independent transcription of the phage neck fiber (*nf*) genes. Expression of the *nf* genes located adjacent to the *phoD* locus in A-Lf14, A-Alj1 and A-Hlh1 during infection. In all three phages, the transcription profiles of the *nf* genes were distinct from each other and from the adjacent *phoD* gene, indicating they are not co-transcribed as part of an operon. (**e**) Expression of the homologous *nf* gene in A-1(L). Transcript levels of the *nf* genes in A-1(L) during infection under +P conditions. (**f**) Concurrent transcription of an overlapping gene pair in A-1(L) and *Anabaena* sp. PCC 7120. Expression of the forward-oriented *tnpB*_fwd_ gene and the novel reverse-oriented *tnpB*_rev_ open reading frame during A-1(L) (which carries a phage-encoded *tnpB* copy) and with A-Lf14, A-Alj1 and A-Hlh1 (which lack *tnpB*) infection, confirming both strands of this locus are transcribed, with strong induction observed specifically during A-1(L) infection. The numbers in panels **d**, **e** and **f** indicate the respective gene cluster numbers (Supplementary Table 4). Data are presented as mean ± SD (n ≥ 3). Statistical significance was determined by Student’s t-test; ns, not significant; *P < 0.05, **P < 0.01, ***P < 0.001, ****P < 0.0001.

Conversely, the *tnpB* gene was uniquely retained in A-1(L) and lacking in the novel isolates. TnpB is a compact, RNA-guided DNA endonuclease ancestrally related to CRISPR-Cas systems, often involved in transposon maintenance and site-specific DNA cleavage^38–40^. The phage’s *tnpB* sequence is identical to five gene copies present in the host *Anabaena* sp. PCC 7120 chromosome (*all3986*, *alr4104*) and its alpha and gamma plasmids *(alr7228*, *alr7231*, *all8010*), indicating recent acquisition or exchange with host mobile genetic elements^42^. Previous studies have shown that these endogenous insertion sequences (ISs) frequently disrupt host genes; for instance, the IS elements carrying *all3986* and *alr4104* each split a two-component histidine kinase family gene, while *alr7231* interrupts a Ser/Thr kinase family gene^42^. This pattern suggests that IS-associated *tnpB* copies are active mobile elements capable of genome rearrangement. Significantly, we found that the *tnpB* locus exhibits a remarkable dual-strand coding architecture: the forward strand encodes the TnpB protein (TnpB_fwd_), while the reverse strand features a long reading frame, potentially encoding a previously undescribed 347-AA protein (TnpB_rev_). Identification of MSTVRASMSSCTQR peptides matching the first 14 amino acids of TnpB_rev_ in two different publicly available proteomics datasets from *Anabaena* sp. PCC 7120^66^ indicates that TnpB_rev_ is produced. BLASTP analysis revealed that the predicted TnpB_rev_ protein shares limited similarity solely with a hypothetical protein (NSP_9610) from *Nodularia spumigena* CCY9414 (52% identity, 67% similarity in an 81% overlap), highlighting its highly novel and lineage-specific origin.

Transcriptional analysis revealed that both forward-strand (*tnpB*_fwd_) and reverse-strand (*tnpB*_rev_) open reading frames are transcribed in uninfected *Anabaena* sp. PCC 7120 (Fig. 6f). Upon infection with A-1(L), which carries a phage-encoded *tnpB* copy, expression of both *tnpB*_fwd_ and *tnpB*_rev_ increased dramatically (Fig. 6f), suggesting that transcriptional induction primarily originated from the phage-borne *tnpB* locus. In contrast, infection with A-Lf14, A-Alj1, and A-Hlh1, all of which lacks *tnpB,* only a modest upregulation of both *tnpB*_fwd_ and *tnpB*_rev_ (Fig. 6f). This result suggests that host-encoded *tnpB* copies are not strongly induced by viral infection in the absence of a phage-borne copy. Nonetheless, the observed basal or slightly elevated transcription could still support a functional role for host *tnpB* in responding to viral challenge, although its contribution may be distinct from that of the phage-encoded system.

We hypothesize that the TnpB system in phage A-1(L) as well as the host-encoded copies, may play a role in phage genome isomer formation and site-specific recombination. The invertible genomic elements present in these phages require precise DNA cleavage and recombination to switch between alternate states, a process that can influence host tropism. An RNA-guided endonuclease like TnpB, potentially directed by a phage-encoded non-coding RNA analog, could facilitate such targeted DNA rearrangements. Alternatively, TnpB could function in a manner analogous to its role in transposon “homing”, cleaving at donor sites to promote homologous recombination that ensures retention of the phage element within host replicons, possibly aiding in lysogeny or genome integration. The co-expression of the novel reverse-strand protein TnpB_rev_ adds another layer of complexity, potentially acting as a regulator, inhibitor, or partner in a multi-protein complex modulating TnpB_fwd_ activity or DNA repair pathways.

## Conclusion

This study elucidates the multifaceted adaptation mechanisms of *Caudoviricetes* cyanophages infecting filamentous nitrogen-fixing cyanobacteria. Employing super-resolution microscopy and cryo-SEM, we visualized infection progression in living filaments with unprecedented detail, revealing striking heterogeneity in infection outcomes even among adjacent cells of the same filament, as well as potential cell-to-cell propagation of virions in later stages. The isolation of three novel phages (A-Lf14, A-Alj1 and A-Hlh1) expands the experimental toolkit for dissecting virus-host interactions in *Anabaena* sp. PCC 7120, a model organism for nitrogen fixation and cellular differentiation. Complementing these phenotypic insights, a central observation is the identification of a large invertible genomic region that co-exists as two stable isomers within phage populations. This genomic isomerism, independent of phylogenetic relatedness, together with the observed infection heterogeneity, represents a previously unrecognized form of plasticity that may modulate gene expression or host interaction traits, adding phenotypic diversity to clonal viral stocks. Such multi-level flexibility, spanning from cellular infection patterns to genome architecture, likely enhances adaptive potential in fluctuating environments, a critical advantage for phages infecting metabolically plastic cyanobacteria.

Our results highlight that cyanophage infection, replication and lysis dynamics are not binary, but rather are modulated by environmental cues, such as light intensity, nitrogen or phosphorus availability. This responsiveness underscores tight coupling between phage replication and host metabolic state, reflecting the evolutionary pressure to synchronize infection with resource availability. Importantly, comparative genomics revealed striking functional divergence driven by accessory genes: the newly isolated cyanophages acquired a gene encoding PhoD, enabling phosphate scavenging under limitation, while A-1(L) retained a unique dual-coding *tnpB* locus with potential roles in site-specific recombination. These findings illustrate how closely related phages occupy distinct ecological niches through the gain or loss of key metabolic or genome-modifying genes.

Collectively, this work bridges genomic features with environmental adaptation, demonstrating that cyanophages employ multi-layered strategies, including genomic isomerism, accessory gene acquisition, and kinetic plasticity, to navigate host physiological changes and resource fluctuations. These insights advance the broader field of phage ecology by revealing how mobile genetic elements and environmental cues shape virus-host dynamics in freshwater ecosystems. Future research focusing on the regulatory mechanisms of genomic isomer switching and the biochemical function of TnpB will further unravel the evolutionary strategies of cyanophages, with implications for understanding microbial community structure and biogeochemical cycling in aquatic environments.

## Methods

### Cultivation of cyanobacterial strains

Cyanobacterial strains were maintained on BG-11 agar plates. A single colony was inoculated into 20 mL of liquid BG-11 medium and grown under continuous white light at an intensity of 50 μmol photons m^-2^ s^-1^, at 30°C with shaking at 150 rpm. Cultures were incubated for 6–7 days until they reached the exponential growth phase (OD_730_ = 0.5–0.8).

### Cyanophage isolation and purification

Planktonic and benthic samples were collected from 36 sampling sites across freshwater ecosystems, including lakes, ponds, and wetlands, in Heilongjiang Province, northeastern China, during 2022–2023. Planktonic material was harvested via horizontal towing at a depth of 0.5 m using 20-μm mesh nets. Benthic specimens were collected from substrates by scraping with sterile brushes or forceps. All samples were immediately transported on ice and stored at −80 °C until processing. For sample preparation, solid specimens were suspended in 500 μ of sterile water. The samples from certain site were homogenized, clarified by gravitational sedimentation, and then filtered through 0.22-μm membranes (Merck Millipore) to obtain cell-free environmental filtrates.

For primary infection assays, thirty-six aliquots of exponentially growing *Anabaena* sp. PCC 7120 were prepared by resuspending cultures in 36 mL BG-11 medium to an OD_730_ of 0.2. Each aliquot was inoculated with 100 μ of environmental filtrate from one of the sampling sites in 24-well plates (900 μ /well). Cultures exhibiting complete lysis within 24–48 hours, evidenced by chlorosis and culture clarification, were centrifuged at 12,000 × g for 10 minutes. The supernatants were then filtered through 0.22-μm membranes to obtain primary cyanophage suspensions. This process of centrifugation, filtration, and reinfection was repeated for three successive rounds to enrich the cyanophage population. Ultimately, three independent cyanophage strains were isolated and designated A-Lf14, A-Alj1, and A-Hlh1 for subsequent genomic analysis.

### Cyanophage genome sequencing and annotation

Genomic DNA was extracted from purified cyanophage particles using the UNIQ-10 Column Virus DNA Isolation Kit (Sangon Biotech). DNA quality was assessed by spectrophotometry (OD_260/280_ = 1.8–2.0), and the concentration was quantified to ensure a minimum of 6 μg per sample. Fragment libraries were constructed from high-quality DNA and sequenced on the Illumina NovaSeq 6000 platform (2 × 150 bp; Biozeron Biotechnology).

Raw sequencing reads were processed with Trimmomatic v0.36, which involved quality trimming by removing bases with Q-scores <20 and adapter sequences. The optimal results for the assembly were obtained by splicing the optimized sequence with multiple Kmer parameters using ABySS (http://www.bcgsc.ca/platform/bioinfo/software/abyss). GapCloser (https://sourceforge.net/projects/soapdenovo2/files/GapCloser/) was applied to fill up the remaining local inner gaps and correct single-base polymorphisms, ultimately enhancing the accuracy and completeness of the final assembly.

Open reading frames (ORFs) were predicted using two annotation pipelines. The first pipeline used GeneMarkS v4.32 (http://exon.gatech.edu/GeneMark/genemarks.cgi) with heuristic model selection, and the second employed RAST (http://rast.nmpdr.org/rast. cgi) via the PATRIC platform. Consensus ORF calls were generated by merging overlapping predictions from both pipelines. All primers used for validation of genome sequences and orientation isomers are listed in Supplementary Table 9.

### Transmission electron microscope (TEM)

Purified cyanophages (10 μ) were applied to carbon-coated copper grids and incubated for 10 minutes. Excess liquid was removed using filter paper. The grids were negatively stained with 3% uranyl acetate for 1-3 minutes, followed by removal of the residual stain with filter paper.

Virions were imaged using a JEM-1200EX transmission electron microscope (JEOL, Tokyo, Japan) operating at an acceleration voltage of 80 kV. Morphometric analysis focused on the characteristic icosahedral capsids, contractile tails and neck fibers.

### Laser scanning confocal microscope (LSCM)

*Anabaena* sp. PCC 7120 cultures were infected with cyanophages and incubated for 12 hours under standard growth conditions to allow phage adsorption. Host-phage mixtures were stained with DAPI to a final concentration of 1 ng/μ and incubated for 10 minutes at room temperature in the dark. Samples were then mounted on glass slides. Images were captured using a Zeiss laser scanning confocal microscope (Carl Zeiss AG, German) that featured a ×63 oil-immersion objective lens with a numerical aperture of 1.4. Images were recorded in 16-bit, 2,186 × 2,186-pixel format and each pixel was 24 × 24 nm. Phycobilisome autofluorescence of *Anabaena* sp. PCC 7120 was detected with excitation at 548 nm and emission at 561 nm, while DAPI was imaged with excitation at 401 nm and emission at 422 nm. Microscope control and image acquisition used Zeiss software ZEN 3.12.

### Cryo-scanning electron microscopy (Cryo-SEM)

For cryo-SEM analysis, infected cells collected 4-8 hours post-infection (initial lysis events were observed) were prepared by placing 5 μ aliquots between gold-plated copper planchettes (3 mm diameter), followed by plunge-freezing in liquid ethane. Samples were then transferred to a pre-cooled stage (−145 °C) in a Zeiss Ultra Plus HR-SEM (Carl Zeiss AG, Germany) equipped with Gemini optics. Imaging was performed at a beam acceleration voltage (BAV) of 2 kV and a working distance (WD) of 4.9 or 7 mm, using in-lens secondary electron detectors for signal collection.

### Plaque assay for cyanophage quantification

Cyanophage titers were determined using the double-agar overlay method. Exponentially growing *Anabaena* sp. PCC 7120 cells were harvested at OD_730_ of 0.5 and mixed with molten BG-11 soft agar (0.6%) at 45 °C. Aliquots (7 mL) containing 10^8^ cells/mL were overlaid onto BG-11 hard agar plates (1.5% agar). Serial dilutions of cyanophage samples (tenfold dilutions in BG-11 medium) were combined with 700 μ of host cell suspension. After a 15-min adsorption period at 30 °C, 100 μ of each mixture was plated in triplicate. Plates were incubated at 30 °C under continuous illumination of 50 μmol photons m^-2^ s^-1^ for 72 hours.

Plaques were enumerated when clearly visible, typically after 3–4 days. Viral titers were calculated using the following formula with three technical triplicates per dilution. PFU/mL = (plaque count × dilution factor) / inoculum volume.

### Host Specificity Analysis

Eleven cyanobacterial strains (*Anabaena* sp. PCC 7120, *Anabaena variabilis* FACHB-319, *Nostoc punctiforme* ATCC 29133, *Nostoc* sp. NIES-2110, *Nostoc calcicola* FACHB 389, *Nostoc spongiaeforme* FACHB 130, *Scytonema* sp. NIES-2130, *Nostoc* ZY3-15, *Nostoc* ZY3-38, *Trichormus* sp. ZY3-43, *Limnospira platensis* MMECC-001, *Synechococcus elongatus* PCC 7942), primarily from the genus *Nostoc* and maintained in our laboratory, along with *Anabaena* sp. PCC 7120, were cultured under standard conditions. Cells were then harvested and resuspended in fresh BG-11 medium to an OD_730_ of 0.2.

Infection assays were performed in 24-well plates, with each well containing 1 mL of cyanobacterial suspension inoculated with 10 μ of cyanophage lysate, resulting in a MOI of 5. Plates were incubated at 30 °C with orbital shaking at 100 rpm under identical light conditions.

Lytic activity was assessed after 24–48 hours by quantifying chlorophyll degradation and confirming cell lysis via microscopy. Strains exhibiting a >50% reduction in OD_730_ relative to uninfected controls were classified as susceptible.

### Infection Kinetics Under Environmental Stress

*Anabaena* sp. PCC 7120 cultures were adjusted to OD_730_ of 0.8 in BG-11 medium. For light/dark assays, 200 mL cultures were infected with cyanophages at multiplicity of infection 0.1 and divided into duplicate flasks: one maintained under standard illumination (50 μmol photons m^-2^ s^-1^) and another in complete darkness, both at 30°C. For nitrogen response assays, parallel cultures were prepared in nitrogen-replete BG-11 and nitrogen-depleted BG-11 media. Aliquots (20 mL) were infected identically and incubated under standard light.

For phosphorus response assays, parallel cultures were prepared in phosphorus-replete BG-11 and phosphorus-depleted BG-11 media. Aliquots (20 mL) were infected identically and incubated under standard light.

Across all conditions, 2 mL samples were collected at 12-hour intervals for 48 hours post-infection. Cell density was quantified by OD_730_ measurements. Uninfected controls under each condition served as references.

### Viptree analysis and library creation

The phage library was created by combining known cyanophages from the Virus-Host Database^57^ and GenBank. To avoid unnecessary data duplication, phages with the same sequences were deleted, only one member per phage species was used and only complete genomes were allowed in the library. For the tree figure, the Viptree algorithm was used^56^ and the host information was taken from Virus-Host Database^57^ and GenBank. The taxonomy information of the phages and hosts was taken from NCBI.

### Quantitative real-time PCR

For RNA extraction, 2 m samples were harvested at 6 h post-infection from both infected and uninfected *Anabaena* sp. PCC 7120 cultures and immediately mixed with a pre-chilled lysis buffer to preserve RNA integrity. Total RNA was purified using the Vazyme Bacterial RNA Extraction Kit. Residual genomic DNA was eliminated, and cDNA was synthesized with the iScript III RT Supermix for qPCR (+gDNA wiper) kit (Vazyme) following the manufacturer’s protocol. Quantitative PCR was performed using a SYBR Premix ExTaqTM (Takara, Dalian, China) on a Roche LightCycler® 480 II sequence detection system (Roche, Shanghai, China). The housekeeping gene *rnpB* (encoding RNase P subunit B) of *Anabaena* sp. PCC 7120 served as an internal reference. Primer sequences for target genes (*phoD*, neck fiber genes, and *tnpB*) are provided in Supplementary Table 9. All qRT-PCR assays were carried out in three biological replicates, which yielded consistent results.

### Reporting summary

Further information on research design is available in the Nature Portfolio Reporting Summary linked to this article.

## Data Availability

The genomic sequences of A-Lf14, A-Alj, A-Hlh and A-1(L) have been deposited in GenBank under accession numbers PX432479 to PX432482, respectively. Source data are provided with this paper.

## Acknowledgements

Financial support for this work was provided by the National Key Research and Development Program of China (2021YFA0909700 and 2024YFB4106501), the Joint Sino-German Research Program (M-0214 to WRH and XL), the DFG (HE 2544/14-2 and HE 2544/20-1/2 to WRH), the Strategic Priority Research Program of Chinese Academy of Sciences (XDB1290103), the National Natural Science Foundation of China (32570172), the QIBEBT International Cooperation Project (QIBEBT ICP202301), the Key Research and Development Program of Shandong Province (2025CXPT029), the Qingdao Science and Technology Benefit People Demonstration Guide Special Project (24-1-8-xdny-17-nsh) and the Shandong Taishan Scholarship (to XL). The authors thank Chunpeng Yang, Xiaofei Yan, and Fei Li for their assistance with confocal laser scanning microscopy (CLSM), cryo-scanning electron microscopy (SEM), and transmission electron microscopy (TEM), respectively. We are grateful to Prof. Xudong Xu for providing the cyanophage A-1(L) and to Prof. Yan Liu for providing part of the environmental samples. We also thank Alexander Mitrofanov and Sven Hauns for the algorithmic search for anti-CRISPR genes. We also thank Dr. Zhuo Chen for his help with the proteomics data analysis.

## Author Contributions Statements

WRH, XL and TZ designed the study. YS and TZ isolated and sequenced the cyanophages and conducted the morphological and physiological experiments. MZ did the majority of bioinformatic analyses. VR maintained phage and cyanobacterial stocks and performed expression analyses. YZ and TZ carried out the environmental sampling. YS, MZ, TZ and WRH wrote the manuscript with contributions from all authors.

## References

1 Zhou, Z. et al. Unravelling viral ecology and evolution over 20 years in a freshwater lake. Nat Microbiol 10, 231–245 (2025).

2 Zimmerman, A. E. et al. Metabolic and biogeochemical consequences of viral infection in aquatic ecosystems. Nat. Rev. Microbiol. 18, 21–34 (2020).

3 Li, Q., Yang, F. & Zhou, C. Z. Cyanophages: Billions of years of coevolution with cyanobacteria. Annu. Rev. Microbiol. 79, 639–661 (2025).

4 Nair, S. et al. Engineering microbiomes to enhance macroalgal health, biomass yield, and carbon sequestration. Green Carbon 3, 63–73 (2025).

5 Golden, J. W. & Yoon, H.-S. Heterocyst development in *Anabaena*. Curr. Opin. Microbiol. 6, 557–563 (2003).

6 Nieves-Morión, M. et al. Single-cell measurements of fixation and intercellular exchange of C and N in the filaments of the heterocyst-forming cyanobacterium *Anabaena* sp. strain PCC 7120. mBio 12, e01314–01321 (2021).

7 Kolan, D. et al. Tradeoffs between phage resistance and nitrogen fixation drive the evolution of genes essential for cyanobacterial heterocyst functionality. ISME J 18 (2024).

8 Higazi, M. et al. Nitrogen availability shapes evolution of phage resistance in cyanobacteria. ISME J 19 (2025).

9 Adolph, K. W. & Haselkorn, R. Isolation and characterization of a virus infecting the blue-green alga *Nostoc muscorum*. Virology 46, 200–208 (1971).

10 Koz’iakov, S., Gromov, B. V. & Khudiakov, I. A-1(L)-cyanophage of the blue-green alga *Anabaena variabilis*. Mikrobiologiia 41, 555–559 (1972).

11 Hu, N. T., Thiel, T., Giddings, T. H., Jr. & Wolk, C. P. New *Anabaena* and *Nostoc* cyanophages from sewage settling ponds. Virology 114, 236–246 (1981).

12 Chénard, C. et al. Viruses infecting a freshwater filamentous cyanobacterium ( *Nostoc* sp.) encode a functional CRISPR array and a proteobacterial DNA polymerase B. mBio 7, e00667–00616 (2016).

13 Yu, R. C. et al. Structure of the intact tail machine of *Anabaena* myophage A-1(L). Nat. Commun. 15, 2654 (2024).

14 Khudiakov, I. & Gromov, B. V. Temperate cyanophage A-4 (L) of the blue-green alga *Anabaena variabilis*. Mikrobiologiia 42, 904–907 (1973).

15 Padan, E. & Shilo, M. Cyanophages-viruses attacking blue-green algae. Bacteriol. Rev. 37, 343–370 (1973).

16 Patel, R. et al. Complete genome sequence of the *Anabaena* myophage Elbi. Microbiol. Resour. Announc. 10, e0055221 (2021).

17 Zhang, S. et al. Host cyanobacteria killing by novel lytic cyanophage YongM: A protein profiling analysis. Microorganisms 10, 257 (2022).

18 Koonin, E. V. et al. Global organization and proposed megataxonomy of the virus world. Microbiol. Mol. Biol. Rev. 84, e00061–00019 (2020).

19 Turner, D., Kropinski, A. M. & Adriaenssens, E. M. A roadmap for genome-based phage taxonomy. Viruses 13, 506 (2021).

20 Gulyaeva, A. et al. Diversity and ecology of *Caudoviricetes* phages with genome terminal repeats in fecal metagenomes from four dutch cohorts. Viruses 14, 2305 (2022).

21 Luque, D. & Castón, J. R. Cryo-electron microscopy for the study of virus assembly. Nat. Chem. Biol. 16, 231–239 (2020).

22 Ortiz de Ora, L., et al. Phollow reveals in situ phage transmission dynamics in the zebrafish gut microbiome at single-virion resolution. Nat Microbiol 10, 1067–1083 (2025).

23 Dai, W. et al. Visualizing virus assembly intermediates inside marine cyanobacteria. Nature 502, 707–710 (2013).

24 Xiong, Z., Wang, Y., Dong, Y., Zhang, Q. & Xu, X. Cyanophage A-1(L) adsorbs to lipopolysaccharides of *Anabaena* sp. strain PCC 7120 via the tail protein lipopolysaccharide-interacting protein (ORF36). J. Bacteriol. 201, 10.1128/jb.00516–00518 (2019).

25 Howe, M. M. The invertible G segment of phage Mu. Cell 21, 605–606 (1980).

26 van de Putte, P., Cramer, S. & Giphart-Gassler, M. Invertible DNA determines host specificity of bacteriophage Mu. Nature 286, 218–222 (1980).

27 Giphart-Gassler, M., Plasterk, R. H. A. & van de Putte, P. G inversion in bacteriophage Mu: a novel way of gene splicing. Nature 297, 339–342 (1982).

28 Plasterk, R. H., Brinkman, A. & van de Putte, P. DNA inversions in the chromosome of *Escherichia coli* and in bacteriophage Mu: Relationship to other site-specific recombination systems. Proc. Natl. Acad. Sci. U. S. A. 80, 5355–5358 (1983).

29 Kamp, D. et al. Comparative analysis of invertible DNA in phage genomes. Cold Spring Harb. Symp. Quant. Biol. 49, 301–311 (1984).

30 Van de Putte, P. & Goosen, N. DNA inversions in phages and bacteria. Trends Genet. 8, 457–462 (1992).

31 Johnson, R. C., Rice, P. & Craig, N. Site-specific DNA inversion by serine recombinases. *Microbiol*. Spectrum 3, 1–36 (2015).

32 Kennedy, K. E. et al. Genome fusion mediated by the site specific DNA inversion system of bacteriophage P1. Mol. Genet. Genomics 189, 413–421 (1983).

33 Iida, S., Hiestand-Nauer, R., Meyer, J. & Arber, W. Crossover sites cix for inversion of the invertible DNA segment C on the bacteriophage P7 genome. Virology 143, 347–351 (1985).

34 Sandmeier, H., Iida, S. & Arber, W. DNA inversion regions Min of plasmid p15B and Cin of bacteriophage P1: evolution of bacteriophage tail fiber genes. J. Bacteriol. 174, 3936–3944 (1992).

35 Dorgai, L., Oberto, J. & Weisberg, R. A. Xis and Fis proteins prevent site-specific DNA inversion in lysogens of phage HK022. J. Bacteriol. 175, 693–700 (1993).

36 Volkert, F. C. & Broach, J. R. Site-specific recombination promotes plasmid amplification in yeast. Cell 46, 541–550 (1986).

37 Chan, K.-M., Liu, Y.-T., Ma, C.-H., Jayaram, M. & Sau, S. The 2 micron plasmid of *Saccharomyces cerevisiae*: A miniaturized selfish genome with optimized functional competence. Plasmid 70, 2–17 (2013).

38 Karvelis, T. et al. Transposon-associated TnpB is a programmable RNA-guided DNA endonuclease. Nature 599, 692–696 (2021).

39 Nakagawa, R. et al. Cryo-EM structure of the transposon-associated TnpB enzyme. Nature 616, 390–397 (2023).

40 Sasnauskas, G. et al. TnpB structure reveals minimal functional core of Cas12 nuclease family. Nature 616, 384–389 (2023).

41 Faure, G. et al. CRISPR-Cas in mobile genetic elements: counter-defence and beyond. Nat. Rev. Microbiol. 17, 513–525 (2019).

42 Wolk, C. P., Lechno-Yossef, S. & Jäger, K. M. The insertion sequences of *Anabaena* sp. strain PCC 7120 and their effects on its open reading frames. J. Bacteriol. 192, 5289–5303 (2010).

43 Padan, E., Ginzburg, D. & Shilo, M. The reproductive cycle of cyanophage LPP1-G in *Plectonema boryanum* and its dependence on photosynthetic and respiratory systems. Virology 40, 514–521 (1970).

44 Lin, X., Ding, H. & Zeng, Q. Transcriptomic response during phage infection of a marine cyanobacterium under phosphorus-limited conditions. Environ. Microbiol. 18, 450–460 (2016).

45 Waldbauer, J. R. et al. Nitrogen sourcing during viral infection of marine cyanobacteria. Proc. Natl. Acad. Sci. U. S. A. 116, 15590–15595 (2019).

46 Liu, Z. & Wu, C. Response of alkaline phosphatases in the cyanobacterium Anabaena sp. FACHB 709 to inorganic phosphate starvation. Curr. Microbiol. 64, 524–529 (2012).

47 Lin, W., Zhao, D. & Luo, J. Distribution of alkaline phosphatase genes in cyanobacteria and the role of alkaline phosphatase on the acquisition of phosphorus from dissolved organic phosphorus for cyanobacterial growth. J. Appl. Phycol. 30, 839–850 (2018).

48 Keroack, C. D. et al. Comparative chemical genomics in Babesia species identifies the alkaline phosphatase PhoD as a determinant of antiparasitic resistance. Proc. Natl. Acad. Sci. U. S. A. 121, e2312987121 (2024).

49 Kageyama, H. et al. An alkaline phosphatase/phosphodiesterase, PhoD, induced by salt stress and secreted out of the cells of Aphanothece halophytica, a halotolerant cyanobacterium. Appl. Environ. Microbiol. 77, 5178–5183 (2011).

50 Luo, H., Benner, R., Long, R. A. & Hu, J. Subcellular localization of marine bacterial alkaline phosphatases. Proc. Natl. Acad. Sci. U. S. A. 106, 21219–21223 (2009).

51 Wang, X. et al. Narrow distribution of cyanophage *psbA* genes observed in two paddy waters of Northeast China by an incubation experiment. Virol. Sin. 31, 188–191 (2016).

52 Wang, X. et al. Novel groups and unique distribution of phage *phoH* genes in paddy waters in Northeast China. Sci. Rep. 6, 38428 (2016).

53 Li, X., Sun, Y., Liu, J., Yao, Q. & Wang, G. Molecular diversity of Cyanopodoviruses in two coastal wetlands in Northeast China. Curr. Microbiol. 76, 863–871 (2019).

54 Rippka, R., Deruelles, J., Waterbury, J. B., Herdman, M. & Stanier, R. Y. Generic Assignments, Strain Histories and Properties of Pure Cultures of Cyanobacteria. J. Gen. Microbiol. 111, 1–61 (1979).

55 Ou, T., Liao, X.-Y., Gao, X.-C., Xu, X.-D. & Zhang, Q.-Y. Unraveling the genome structure of cyanobacterial podovirus A-4L with long direct terminal repeats. Virus Res. 203, 4–9 (2015).

56 Nishimura, Y. et al. ViPTree: the viral proteomic tree server. Bioinformatics 33, 2379–2380 (2017).

57 Mihara, T. et al. Linking virus genomes with host taxonomy. Viruses 8, 66 (2016).

58 Zhu, J. et al. Phylogenomics of five *Pseudanabaena* cyanophages and evolutionary traces of horizontal gene transfer. *Environ*. Microbiome 18, 3 (2023).

59 Wandera, K. G. et al. Anti-CRISPR prediction using deep learning reveals an inhibitor of Cas13b nucleases. Mol. Cell 82, 2714–2726.e2714 (2022).

60 Bancroft, I. & Smith, R. J. Restriction mapping of genomic DNA from five cyanophages infecting the heterocystous cyanobacteria *Nostoc* and *Anabaena*. New Phytol. 113, 161–166 (1989).

61 Khudyakov, I. & Wolk, C. P. Evidence that the hanA gene coding for HU protein is essential for heterocyst differentiation in, and cyanophage A-4(L) sensitivity of, Anabaena sp. strain PCC 7120. J. Bacteriol. 178, 3572–3577 (1996).

62 Casjens, S. R. & Gilcrease, E. B. Determining DNA packaging strategy by analysis of the termini of the chromosomes in tailed-bacteriophage virions. Methods Mol. Biol. 502, 91–111 (2009).

63 Li, S. et al. Scrutinizing virus genome termini by high-throughput sequencing. PLoS One 9, e85806 (2014).

64 Garneau, J. R., Depardieu, F., Fortier, L.-C., Bikard, D. & Monot, M. PhageTerm: a tool for fast and accurate determination of phage termini and packaging mechanism using next-generation sequencing data. Sci. Rep. 7, 8292 (2017).

65 Nanamiya, H., Tanaka, D., Hiyama, G., Isogai, T. & Watanabe, S. Detection of four isomers of the human cytomegalovirus genome using nanopore long-read sequencing. Virus Genes 60, 377–384 (2024).

66 Zhang, Q., Yu, S., Wang, Q., Yang, M. & Ge, F. Quantitative proteomics reveals the protein regulatory network of Anabaena sp. PCC 7120 under nitrogen deficiency. J. Proteome Res. 20, 3963–3976 (2021).

