## Extended Data Figures 1 to 5 for "Genomic Isomerism and Environmental Adaptation in Cyanophages Infecting Freshwater Cyanobacteria"

**Extended Data Fig. 1**. Three-dimensional localization of phage DNA in *Anabaena* filaments during infection. *Anabaena* filaments infected with cyanophage A-Lf14 were stained with DAPI (pseudocolored green; excitation 401 nm, emission 422 nm) and imaged using a Zeiss LSM 880 confocal microscope equipped with Airyscan super-resolution detection. Images were acquired with a 63× oil-immersion objective (NA 1.4) through a 21-slice Z-stack, followed by joint deconvolution processing to enhance resolution. Phage DNA signals (green) are shown relative to host phycobilisome autofluorescence (pseudocolored magenta; excitation 548 nm, emission 561 nm). The images display two filament morphologies (Infection 1 and Infection 2) captured from the infected culture, illustrating variation within the population. Uninfected control filament (CK) is shown for reference. A video providing a volumetric rendering of the Infection 2 filament is provided in **Supplementary Video 1**.

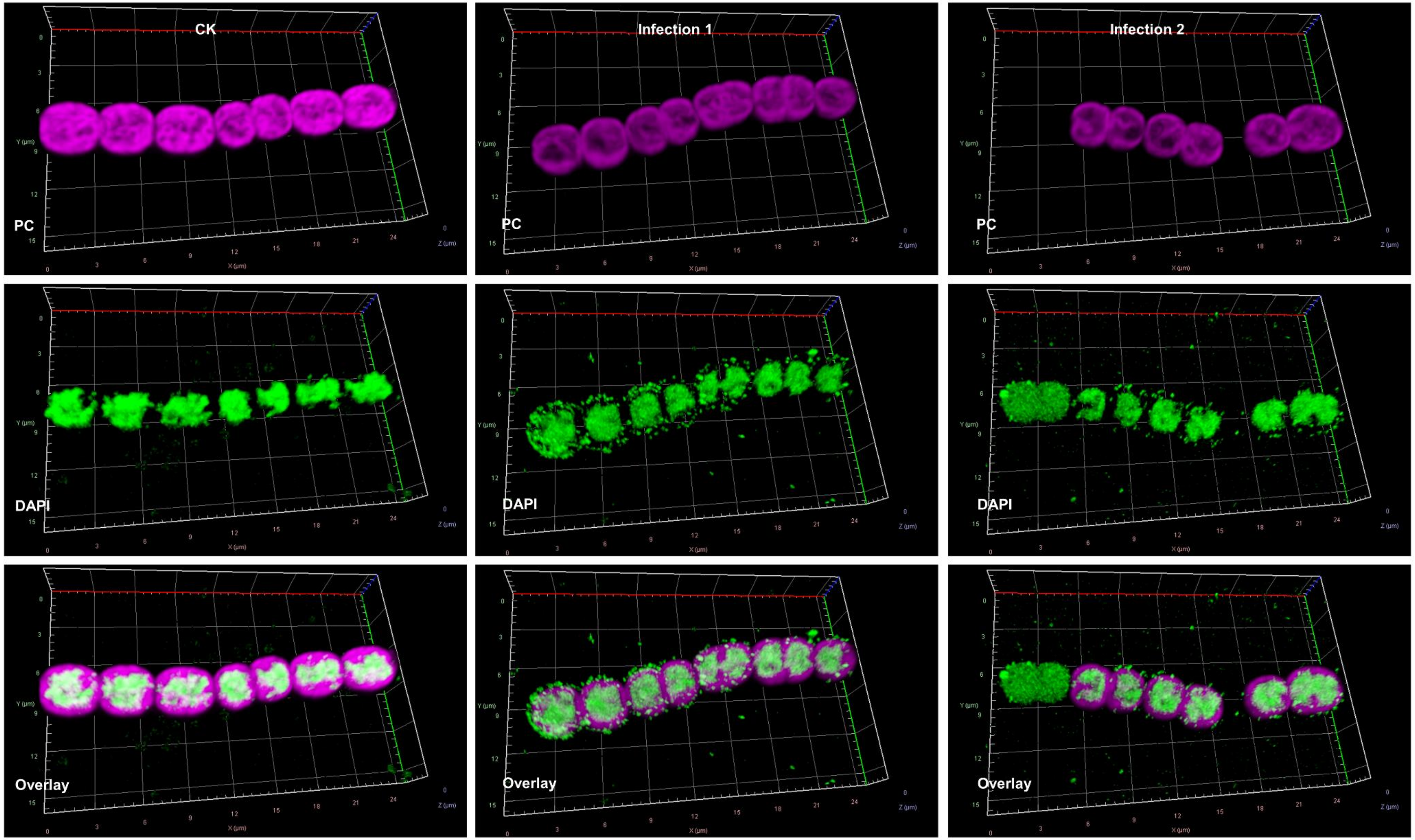

**Extended Data Fig. 2**. Cryo-scanning electron microscopy (cryo-SEM) of *Anabaena* cells at successive stages of phage-induced lysis. *Anabaena* sp. PCC 7120 cells infected with cyanophage A-1(L) were plunge-frozen at 8 h post-infection and imaged by cryo-SEM. (a) A field containing both an uninfected cell (upper left) and a neighboring cell in an advanced stage of lysis (lower right). (b-i) Single-cell views illustrating a progression of phage-induced structural disruption, from early alterations to complete cellular disintegration. Imaging was performed on a Zeiss Ultra Plus HR-SEM operated at 2 kV with a stage temperature of -145 °C.

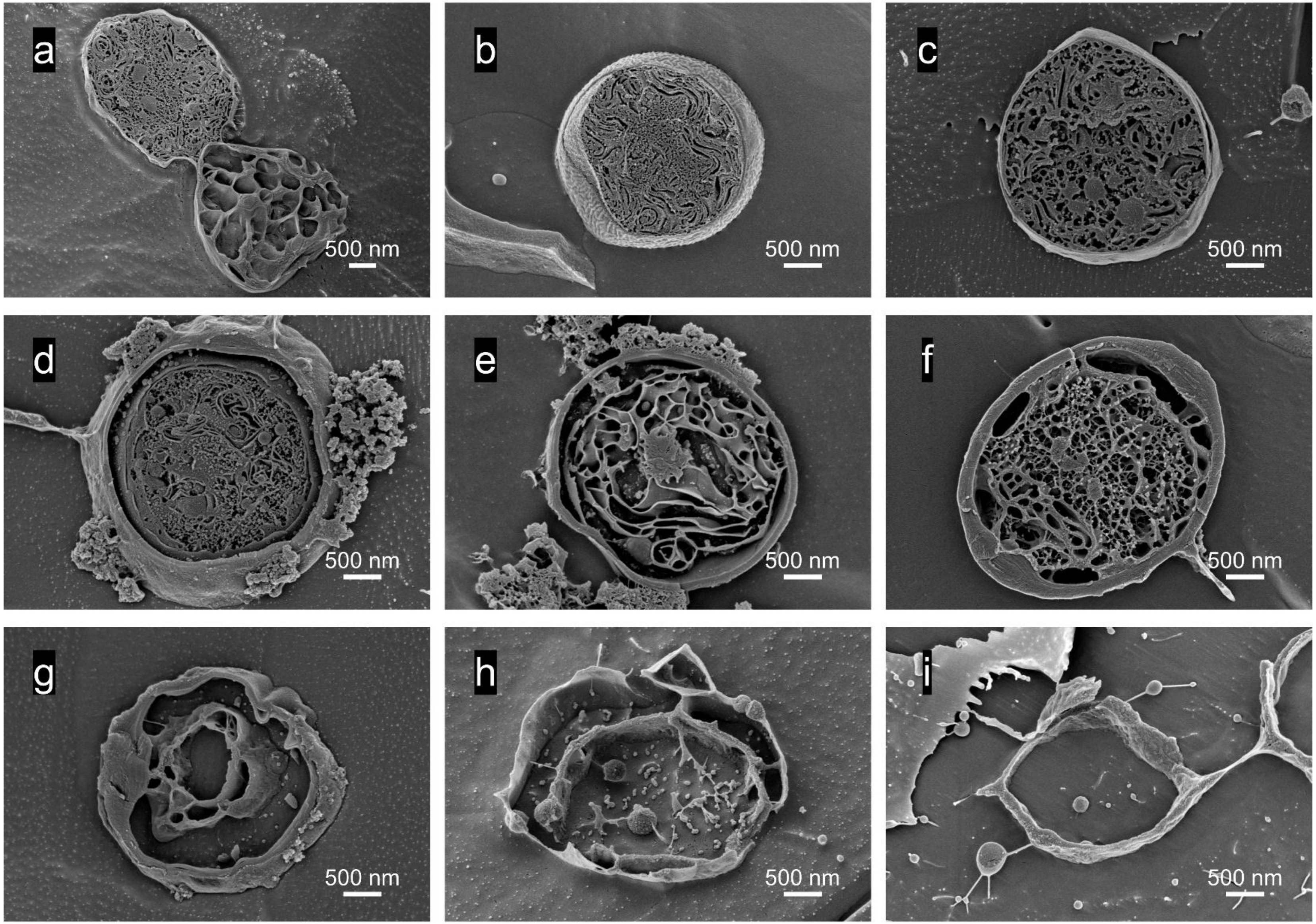

**Extended Data Fig. 3**. Alignment (sequence logo) of IR elements of the nine phage genomes displayed in Fig. 3a.

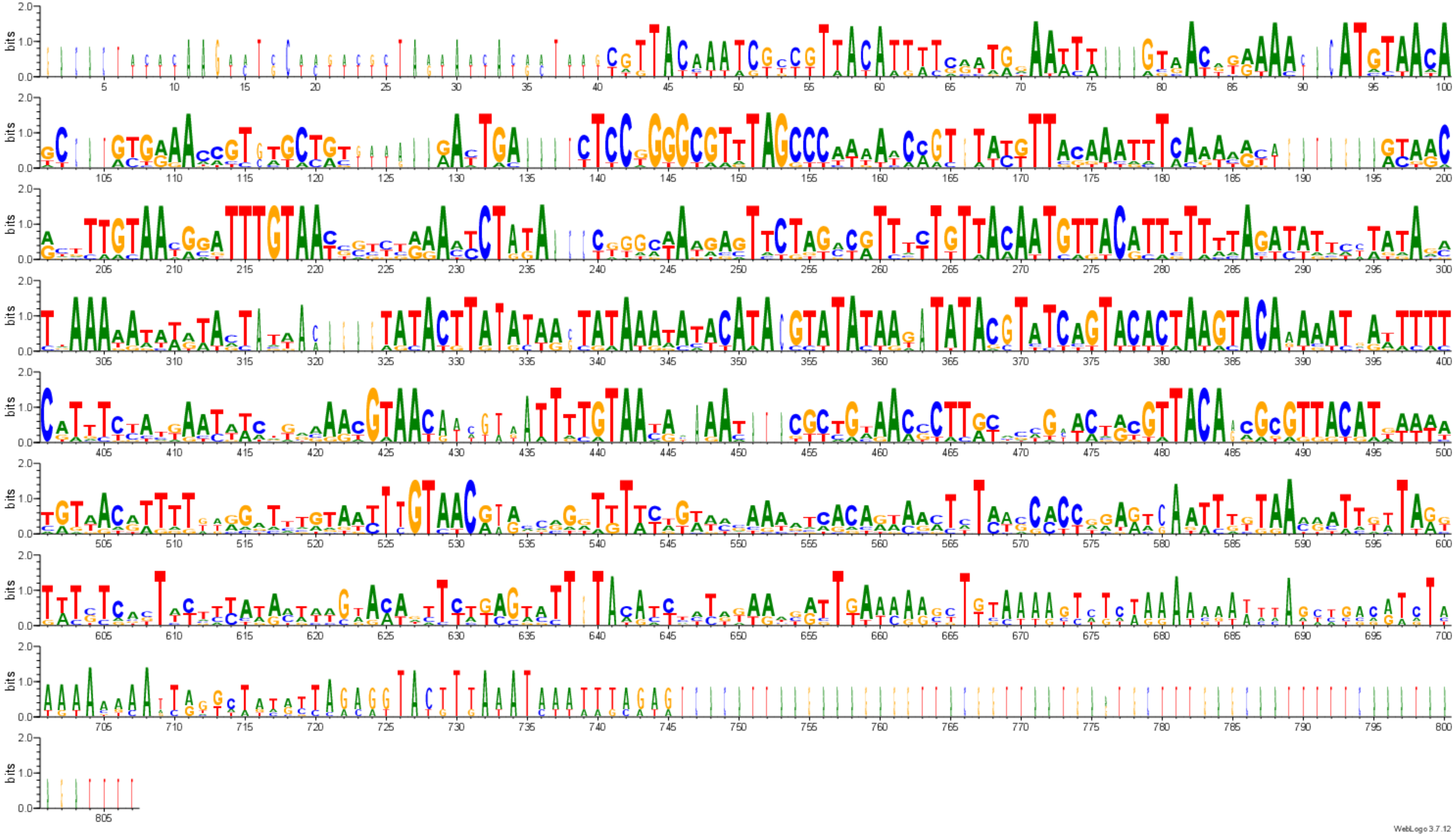

**Extended Data Fig. 4**. Verification of isomeric configurations by Sanger sequencing. PCR products (2.3-4.2 kb) amplified with primers Uni/P1 and Uni/P2 from each cyanophage strain were Sanger sequenced. The assembled contigs were aligned to the corresponding eight reference regions derived from the two isomers of strains A-1(L), A-Lf14, A-Alj1 and A-Hlh1, confirming their nucleotide-level accuracy.

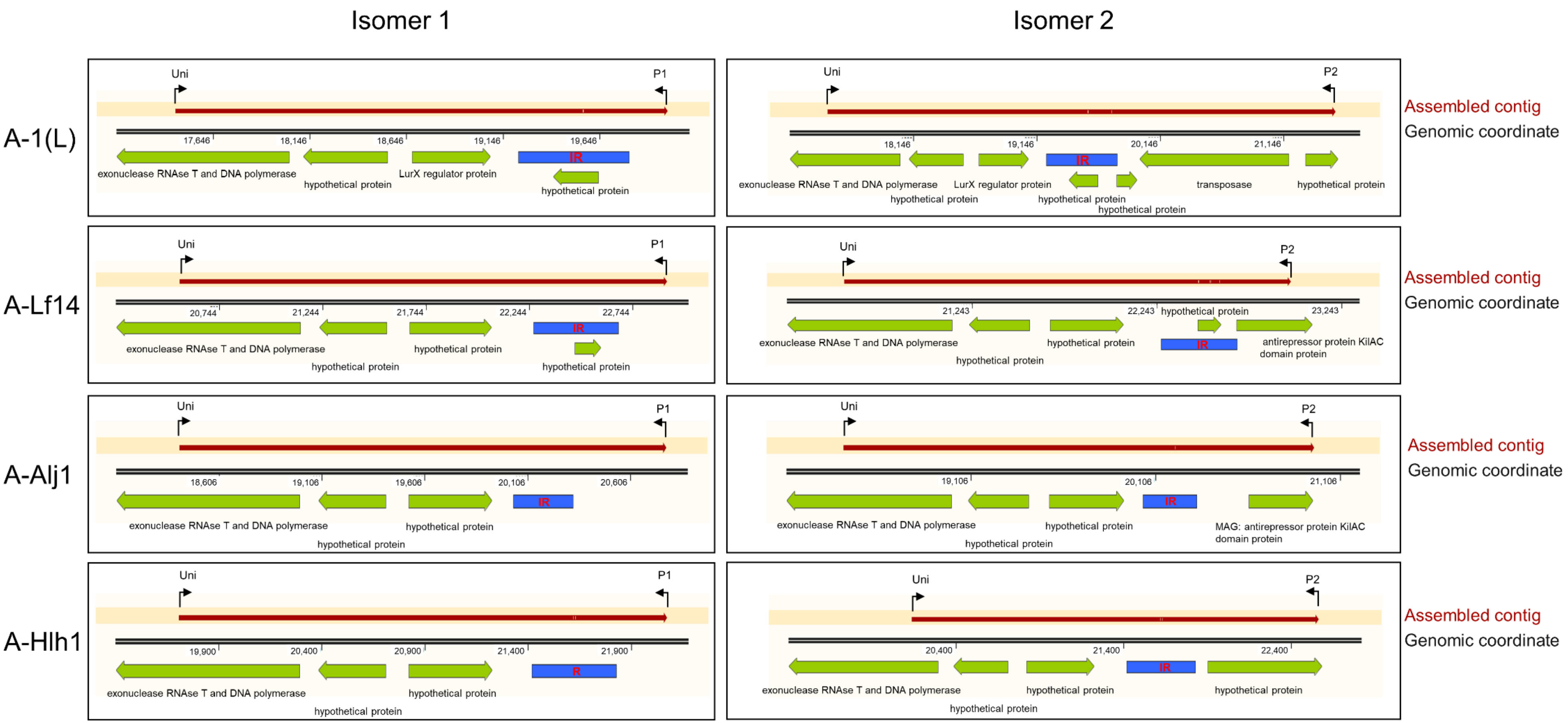

**Extended Data Fig. 5**. Validation of genomic isomers by mapping of sequencing reads. Read mapping of Illumina sequencing data against the two constructed reference genomes for each phage isolate. For isolates where the *de novo* assembly resulted in Isomer 1, the invertible region was reverse-complemented to generate an Isomer 2 reference sequence (and vice versa). The reduced read depth at the invertible region boundaries (indicated by arrows) reflects the repetitive nature of these sequences, which complicates assembly and read mapping and supports the presence of both isomeric configurations in the sequencing library.

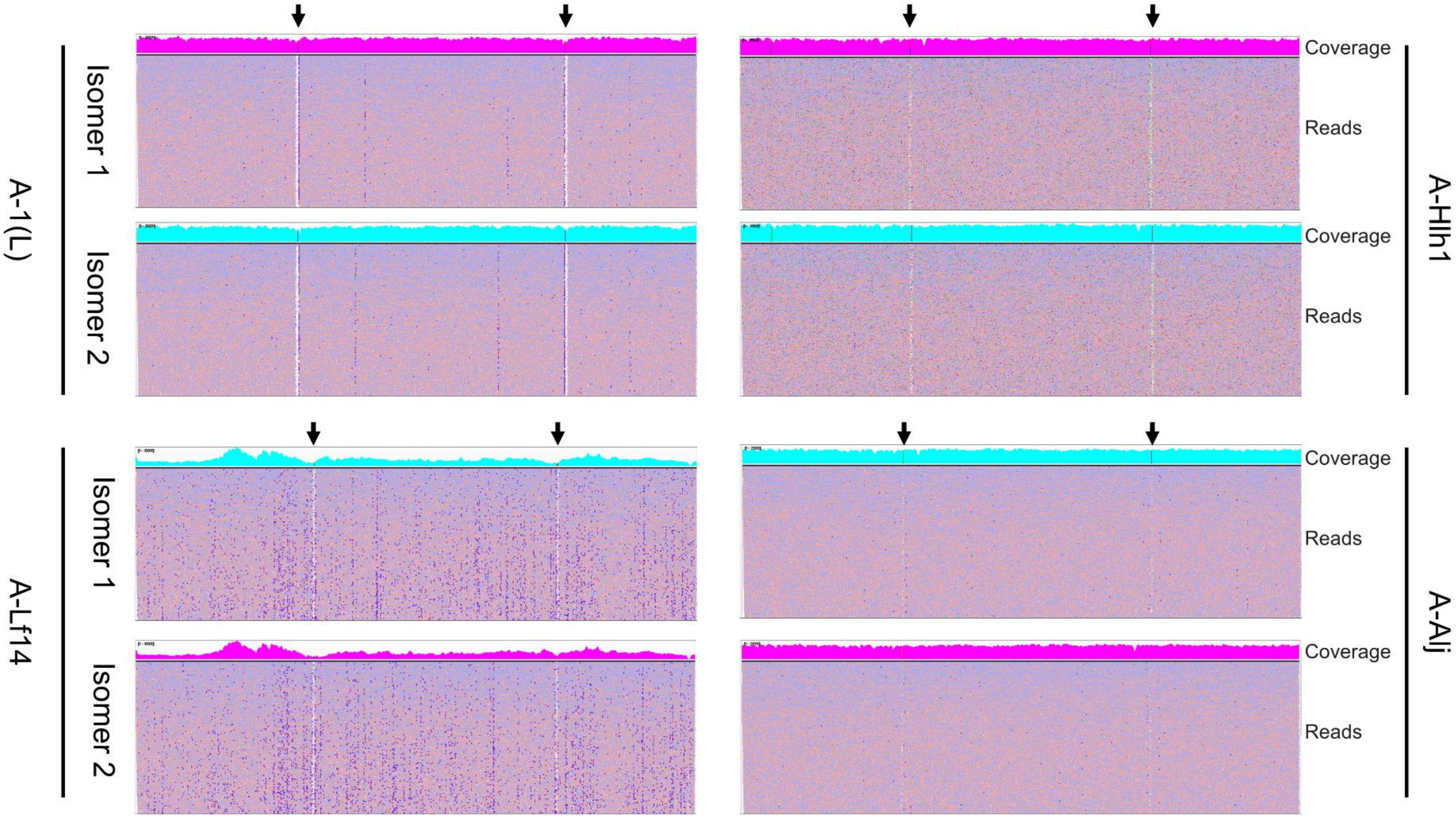
